# Frequent transitions in mating-type locus chromosomal organization in *Malassezia* and early steps in sexual reproduction

**DOI:** 10.1101/2023.03.25.534224

**Authors:** Marco A. Coelho, Giuseppe Ianiri, Márcia David-Palma, Bart Theelen, Rohit Goyal, Aswathy Narayanan, Jeffrey M. Lorch, Kaustuv Sanyal, Teun Boekhout, Joseph Heitman

## Abstract

Fungi in the basidiomycete genus *Malassezia* are the most prevalent eukaryotic microbes resident on the skin of human and other warm-blooded animals and have been implicated in skin diseases and systemic disorders. Analysis of *Malassezia* genomes revealed that key adaptations to the skin microenvironment have a direct genomic basis, and the identification of mating/meiotic genes suggests a capacity to reproduce sexually, even though no sexual cycle has yet been observed. In contrast to other bipolar or tetrapolar basidiomycetes that have either two linked mating-type-determining (*MAT*) loci or two *MAT* loci on separate chromosomes, in *Malassezia* species studied thus far the two *MAT* loci are arranged in a pseudobipolar configuration (linked on the same chromosome but capable of recombining). By incorporating newly generated chromosome-level genome assemblies, and an improved *Malassezia* phylogeny, we infer that the pseudobipolar arrangement was the ancestral state of this group and revealed six independent transitions to tetrapolarity, seemingly driven by centromere fission or translocations in centromere-flanking regions. Additionally, in an approach to uncover a sexual cycle, *Malassezia furfur* strains were engineered to express different *MAT* alleles in the same cell. The resulting strains produce hyphae reminiscent of early steps in sexual development and display upregulation of genes associated with sexual development as well as others encoding lipases and a protease potentially relevant for pathogenesis of the fungus. Our study reveals a previously unseen genomic relocation of mating-type loci in fungi and provides insight towards the discovery of a sexual cycle in *Malassezia*, with possible implications for pathogenicity.

**Significance Statement:** *Malassezia*, the dominant fungal group of the mammalian skin microbiome, is associated with numerous skin disorders. Sexual development and yeast-to-hyphae transitions, governed by genes at two mating-type (*MAT*) loci, are thought to be important for fungal pathogenicity. However, *Malassezia* sexual reproduction has never been observed. Here, we used chromosome-level assemblies and comparative genomics to uncover unforeseen transitions in *MAT* loci organization within *Malassezia*, possibly related with fragility of centromeric-associated regions. Additionally, by expressing different *MAT* alleles in the same cell, we show that *Malassezia* can undergo hyphal development and this phenotype is associated with increased expression of key mating genes along with other genes known to be virulence factors, providing a possible connection between hyphal development, sexual reproduction, and pathogenicity.

## Introduction

The skin microbiome of humans and other animals is composed of a range of organisms, including viruses, bacteria, and fungi. These organisms interact with host skin cells in different ways, including aiding in defense and immunity, inhibiting the growth of opportunistic or pathogenic organisms, and promoting tissue repair and barrier protection (1). Yeasts of the basidiomycete genus *Malassezia* are the dominant fungal residents of the skin of warm-blooded vertebrates and are usually enriched in sebaceous sites (2, 3). This adaptation is linked to their lipid-dependent growth, which is associated with the loss of key genes for fatty acid synthesis (4, 5). At present, the genus *Malassezia* includes 18 formally described species, with *M. restricta*, *M. arunalokei*, *M. globosa*, *M. sympodialis*, and *M. furfur* more commonly observed on human skin regardless of health or disease status (6). While these species are often considered commensals, they can also be mutualistic or pathogenic. Indeed, *Malassezia* has been linked to various skin disorders (3), and recently implicated in Crohn’s disease (7) and progression of colorectal (8) and pancreatic cancer (9). *Malassezia* may also restrict other fungal pathogens, such as *Candida auris*, from colonizing the skin (10). Direct sequencing approaches of diverse environmental sources, such as soil, corals, sponges, deep-sea vents (3, 11), and the skin of healthy human volunteers (12), also revealed that *Malassezia* is more diverse than previously thought.

Whole-genome sequencing of *Malassezia* has provided a wealth of data to explore how this fungal genus has evolved and adapted to the skin microenvironment. Unique genomic features include (i) small and compact genomes (∼7 to 9 Mb) (4, 5); (ii) an increased number of genes encoding lipases, phospholipases, and sphingomyelinases that allow the fungus to exploit abundant extracellular lipids (4, 5); (iii) a low density of repeats and transposable elements possibly associated with loss of the RNA interference machinery (4, 5); (iv) the presence of a dsRNA mycovirus (13, 14); (v) the acquisition of genes via horizontal gene transfer from bacterial lineages commonly found as members of mammalian microbiomes (5, 15); and (vi) karyotypic diversity associated with short centromeres composed of AT-rich sequences (16). These studies also indicated that *Malassezia* species harbor genes necessary to reproduce sexually (5, 17, 18). Despite this, a sexual cycle has yet to be observed in laboratory or natural settings. Due to mechanisms such as recombination and independent assortment during meiosis, sexual reproduction provides a broader range of genetic variation for natural section to act upon, in contrast to asexual reproduction, and has been hypothesized to have a role in fungal pathogenesis (19, 20). In addition, sexual development in basidiomycetous yeast lineages is typically marked by a transition from yeast to hyphae, initiated when cells of compatible mating type fuse. This change in morphology has been linked to virulence in several fungal species (21), and has been well studied in smut fungi that affect cereals and other crops (22).

Mating-type compatibility in heterothallic basidiomycetes is regulated by two mating-type (*MAT*) loci: the pheromone/receptor (*P/R*) locus, which encodes at least one mating pheromone precursor (Mfa) and one pheromone receptor (Pra) involved in mate recognition, and the homeodomain (*HD*) locus that contains two divergently transcribed homeodomain genes, *HD1*/*bE* and *HD2*/*bW*, which control sexual development following cell fusion (23). To mate and successfully produce offspring, cells must carry different, compatible alleles of both *MAT* loci. In most basidiomycetes, the *P/R* and *HD* loci are located on different chromosomes (tetrapolar configuration), but in some species they have become linked, resulting in an expanded chromosomal region that is highly rearranged and suppressed for recombination between opposite mating types (the bipolar configuration) [reviewed in (23, 24)]. Interestingly, the bipolar configuration is found in several basidiomycetes that are associated with animals or plants, either as pathogens or as commensals (e.g., *Cryptococcus* pathogenic species, the barley smut *Ustilago hordei*, and most anther-smut *Microbotryum* species), and it is presumed to have evolved to increase inbreeding (50% chance of compatibility in bipolar compared to 25% in tetrapolar) (23). In contrast, earlier studies of *Malassezia* showed that while the *P/R* and *HD* loci are located on the same chromosome, they are separated by an extended intervening chromosomal region (>100 kb) allowing recombination to still occur to generate novel mating type configurations – an organization known as pseudobipolar (4, 5, 17, 18, 25). Because extant sexual reproduction has yet to be observed for any *Malassezia* species, evidence of recombination is based solely on population genetics studies of natural isolates (17, 26). Furthermore, although *Malassezia* genome sequencing efforts provided an initial view of genome content, most of the available genomes are not assembled as complete chromosomes. Thus, it was unclear if co-localization of the *P/R* and *HD* loci on the same chromosome is shared by all species and ancestral to this group of fungi, or if there are different mating-type loci configurations across species. It was also unknown if the mating-type genes are associated with the yeast-to-hypha transition, which has been sporadically reported for a few *Malassezia* species in association with disease (27).

In this study, we set out to explore the potential for sexual reproduction in *Malassezia* and understand the underlying genetic structures and mechanisms that govern this process. To achieve this, we have assembled and refined the genomes of more than half of the known *Malassezia* species, including a newly identified species isolated from the skin of healthy bats. We performed a series of comprehensive cross-species comparative analyses, illuminating the evolution of telomeres, karyotype, and transitions in *MAT* loci chromosomal structure in this fungal genus. Additionally, we show that *M. furfur* can undergo hyphal development when different mating-type alleles are expressed in the same cell following *Agrobacterium*-mediated transformation. Associated with this morphological transition, we observed upregulation of several genes linked to sexual development along with other genes potentially relevant for pathogenesis. Our findings provide evidence that this remarkable group of fungi have the capacity for mating and sexual reproduction, with a possible link to pathogenicity.

## Results

### Improved genomic resources for the *Malassezia* genus and new insight from telomere-to-telomere assemblies

Of the 18 formally described *Malassezia* species, only six had genomes assembled to completion or near completion (**Fig. 1A**). By combining long- (Oxford nanopore) and short-read (Illumina) sequencing, gapless, telomere-to-telomere, assemblies with high base accuracy were generated for 12 *Malassezia* species, including that of a new species recently isolated from bats (*Malassezia* sp. CBS17886) (28), which will be formally described elsewhere (**Fig. 1A**). To enable comparative genomic analysis, we integrated the available genome assemblies of an additional eight *Malassezia* species, including two strains within the *M. furfur* species complex as they appear to represent closely related but distinct sibling species (26). Hence, the final dataset comprises 20 *Malassezia* species. Based on all assemblies and excluding the genomes of previously identified hybrid strains (26), the *Malassezia* nuclear genomes range from 7.33 Mb (for *M. restricta*) to 9.31 Mb in size (for *Malassezia* sp. CBS17886) and are organized into 6 to 9 chromosomes (**Fig. 1B** and ***SI Appendix*, Table S1**).

**Fig. 1.**
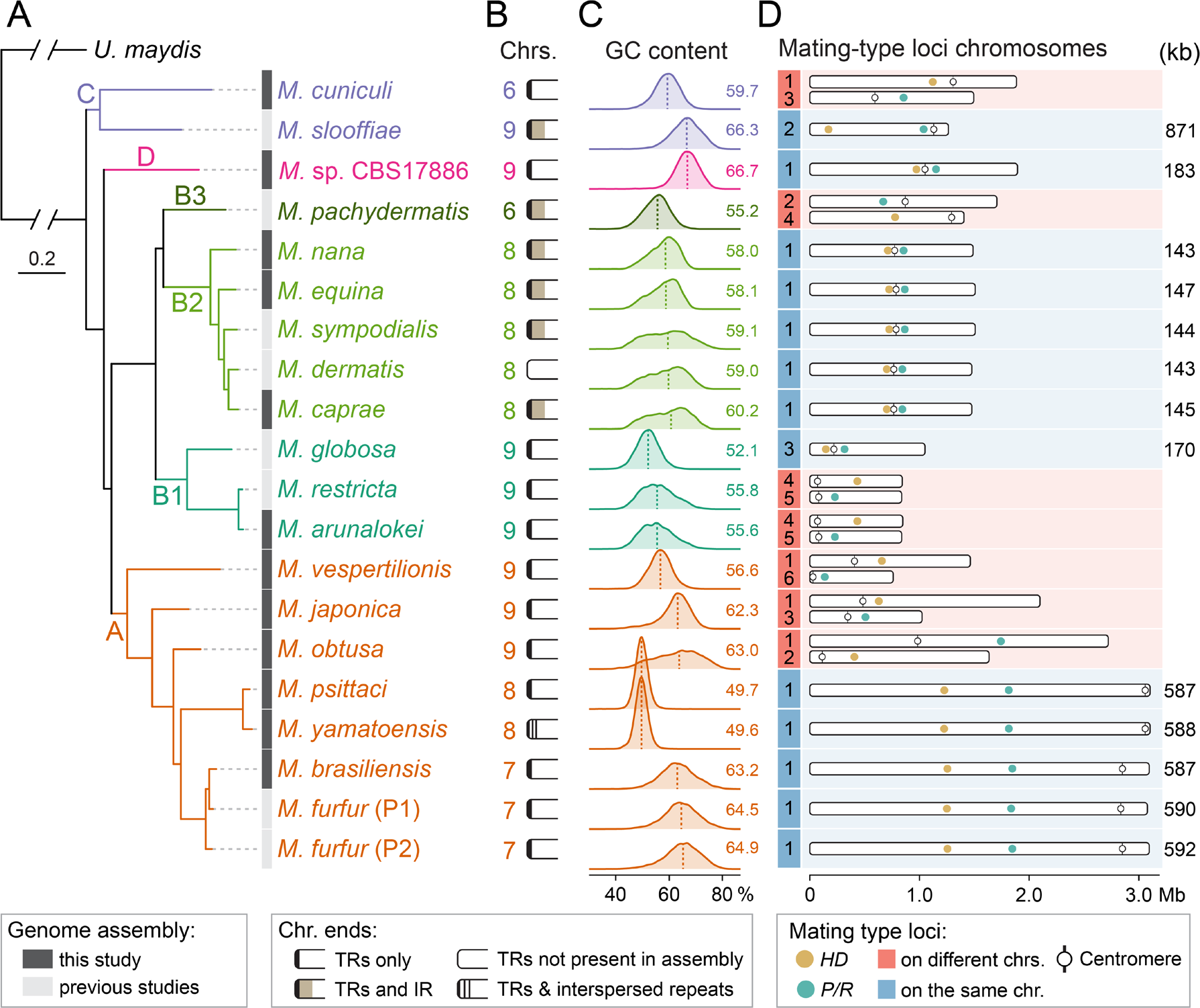
**Phylogenetic diversity, karyotypic variation, and mating-type loci organization in *Malassezia***. (A) Species phylogeny inferred by maximum likelihood analysis using a concatenation-based approach on a data matrix composed of protein alignments of 1,566 single- copy genes shared across all *Malassezia* species and an outgroup (*Ustilago maydis*). All branches are 100% supported (SH-aLRT and UFboot tests; refer to ***SI Appendix,* Fig. S1**). Branch lengths are given in number of substitutions per site (scale bar). Complete genome sequences obtained in this study are marked in dark grey. (B) Chromosome number and structure of the chromosome ends. All assemblies, with the exception of *M. dermatis*, are assembled telomere-to-telomere. In some species, chromosome ends present inverted repeats (IR) beyond the telomeric-repeat sequences (TRs). In *M. yamatoensis*, some of the chromosome ends are expanded and display TRs motifs interspersed with other repeat sequences (refer to ***SI Appendix,* Fig. S2**). (C) Frequency distribution of GC content across species. Mean GC values are shown next to the plot and depicted by vertical dashed lines. Some species present constant GC content across the genome (e.g., *M. yamatoensis*), while others (e.g., *M. obtusa*) have an uneven distribution. (D) Chromosomal localization of the two mating-type loci (pheromone/receptor locus – *P/R*, and homeodomain transcription factor locus – *HD*) with distance between the two loci shown in kb on the right.

To establish the evolutionary relationships within *Malassezia*, gene prediction and annotation were performed, identifying 1,566 single-copy genes shared across all *Malassezia* species and an outgroup (*Ustilago maydis*). Phylogenetic reconstructions based on concatenation and coalescence approaches generated completely congruent and robustly supported trees **(Fig. 1** and ***SI Appendix*, Fig. S1**). This analysis recovered the three known main clades (A, B and C) (5, 12) and confirmed the new species status of *Malassezia* sp. CBS17886 given its placement in a distinct branch (clade D). Clades A and B are sister clades, and clade C is the earliest diverging lineage. Furthermore, an interesting relationship was uncovered between *M. furfur* and *Malassezia brasiliensis*, with *M. furfur* parental lineage 1 (P1) and *M. brasiliensis* being more closely related than the two *M. furfur* parental lineages P1 and P2. This builds upon recent findings proposing that the *M. furfur* complex probably encompasses more than one species and has evolved through a convoluted process involving multiple hybridization events (26).

All *Malassezia* genome assemblies (except *M. dermatis* which was not re-sequenced in this study) had telomeric repeats at the ends of each chromosome. This prompted a closer examination of the telomeric regions, resulting in the recovery of four versions of the telomere-repeat motifs (***SI Appendix*, Table S1**). Apart from *Malassezia pachydermatis* and *Malassezia vespertilionis*, these variants tended to be clade-specific (***SI Appendix*, Table S1**). Upon further examination, distinct species- or clade-specific patterns were observed at the telomere regions. First, in *Malassezia yamatoensis*, 7 of the 16 chromosome ends consist of blocks of telomeric repeats interspersed with unknown repeat sequences (∼650 bp). These blocks can span several kilobases (e.g., ∼42 kb at the left end of chr. 6, ***SI Appendix*, Fig. S2A**) and feature two distinct repeat patterns: a canonical one with multiple tandem copies of the “GGATTA” telomeric repeat motif (right end), and a variant form where the motif “GGATGTC” is intermingled with the telomeric repeat motif (TR, type 1 and 2, respectively, ***SI Appendix*, Fig. S2A**). This unusual telomere structure is not seen in the sister species *Malassezia psittaci* and thus indicates a recent change in *M. yamatoensis* telomere structure. Second, in *Malassezia slooffiae* and in species of sub-clades B2 and B3, inverted repeats are found next to the telomeric repeat tract almost invariably present in all chromosomes (**Fig. 1B**, ***SI Appendix*, Fig. S2B** and **Table S1**). Translation of these sequences revealed a large open reading frame in all species except *M. nana*, each predicted to encode a 1,479- to 1,648-amino-acid protein with a reverse transcriptase domain (PF00078). BLASTP searches of these proteins in GenBank produced top matches to hypothetical proteins of other basidiomycetes, including the *Cryptococcus neoformans* Cnl1 element, a non-long-terminal repeat (non-LTR) retrotransposon found predominantly associated with telomeric repeat sequences (29, 30). In *M. slooffiae*, interstitial telomeric repeats are found between copies of the non-LTR element, but this arrangement is absent in other species (***SI Appendix***, **Fig. S2B**). We thus hypothesize that these elements in *Malassezia* are non-LTR retrotransposons that may target telomere repeats, similar to those reported in other eukaryotes, including fungi (29, 31, 32).

Previous work in *M. sympodialis* and *M. furfur* established that the region with the lowest GC-content on each chromosome coincides with experimentally validated centromeres (16). Accordingly, inspection of the GC content for the new and updated assemblies found a single discrete region with distinctly low GC content on each chromosome, and these regions were assigned as candidate centromeres (***SI Appendix***, **Fig. S3** and **Dataset S1**). This analysis also revealed a broad variation of the GC-content across species, ranging from 49.6% in *M. yamatoensis* to 66.7% in *Malassezia* sp. CBS17886, with most species exhibiting a unimodal GC-content distribution (**Fig. 1C**, and ***SI Appendix*, Fig. S3**). However, six species (*M. sympodialis*, *M. dermatis*, *M. caprae*, *M. nana*, *M. equina*, and *M. obtusa*) displayed an unusual wide GC-content distribution and a partially bimodal profile, suggesting their genomes contain a greater proportion of AT-rich regions (**Fig. 1C**, and ***SI Appendix*, Fig S3**). The biological significance of this irregular GC-content distribution in these species is currently unclear and is planned to be addressed in future studies. Overall, this comprehensive genomic dataset provides an essential resource for further exploring variation across *Malassezia* species and strains, which can now be addressed at gene, chromosomal, and population levels.

### Mating-type loci are conserved across species but exhibit distinct chromosomal organization

Studies in *M. globosa*, *M. sympodialis*, *M. yamatoensis*, and *M. furfur* uncovered a pseudobipolar *MAT* locus configuration with *P/R* and *HD* mating-type loci residing on the same chromosome but separated by large syntenic chromosomal regions (4, 5, 17, 18, 26). Using BLAST searches, we now determined the precise *MAT* loci configurations in all *Malassezia* species. Unexpectedly, out of the 20 species surveyed, 13 have a pseudobipolar configuration, whereas in the remaining seven the *P/R* and *HD* loci are found on different chromosomes (tetrapolar configuration). Notably, both pseudobipolar and tetrapolar configurations were recovered in all of the clades (**Fig. 1D**), which prompted us to delineate the evolutionary trajectory of the *MAT* loci in *Malassezia*.

By performing synteny analyses we first corroborated and expanded previous findings (4, 5, 17, 18, 26) that the pheromone/receptor (*P/R*) locus contains only two divergently oriented genes encoding a pheromone precursor (*MFA*) and a pheromone receptor (*PRA*) and the *HD* locus comprises two divergently transcribed homeodomain genes designated as *bE* (HD1-type) and *bW* (HD2-type), as found in many basidiomycetes (**Fig. 2A**, and ***SI Appendix*, Fig. S4**) (23). Sequence alignment and phylogenetic analysis determined that the *P/R* locus is biallelic (*a1* or *a2*) as the products of the mating pheromone (Mfa1/Mfa2) and pheromone receptor genes (Pra1/Pra2) clustered into two groups (**Figs. 2B** and **2C**). Phylogenetic clustering of alleles by mating type across species, rather than clustering by species, is expected for alleles of genes at the *P/R* locus, as they ceased recombining long ago and have since accumulated substitutions independently (known as trans-specific polymorphism) (33). Further examination of the predicted protein sequences of the mature pheromone found that they were indistinguishable between a few closely related species, namely the Mfa2 of *Malassezia dermatis* vs. *Malassezia nana*, and *M. furfur* (P2) vs. *M. brasiliensis* (**Fig. 2C**). However, it is uncertain whether their respective Mfa1 pheromones are also identical.

**Fig. 2.**
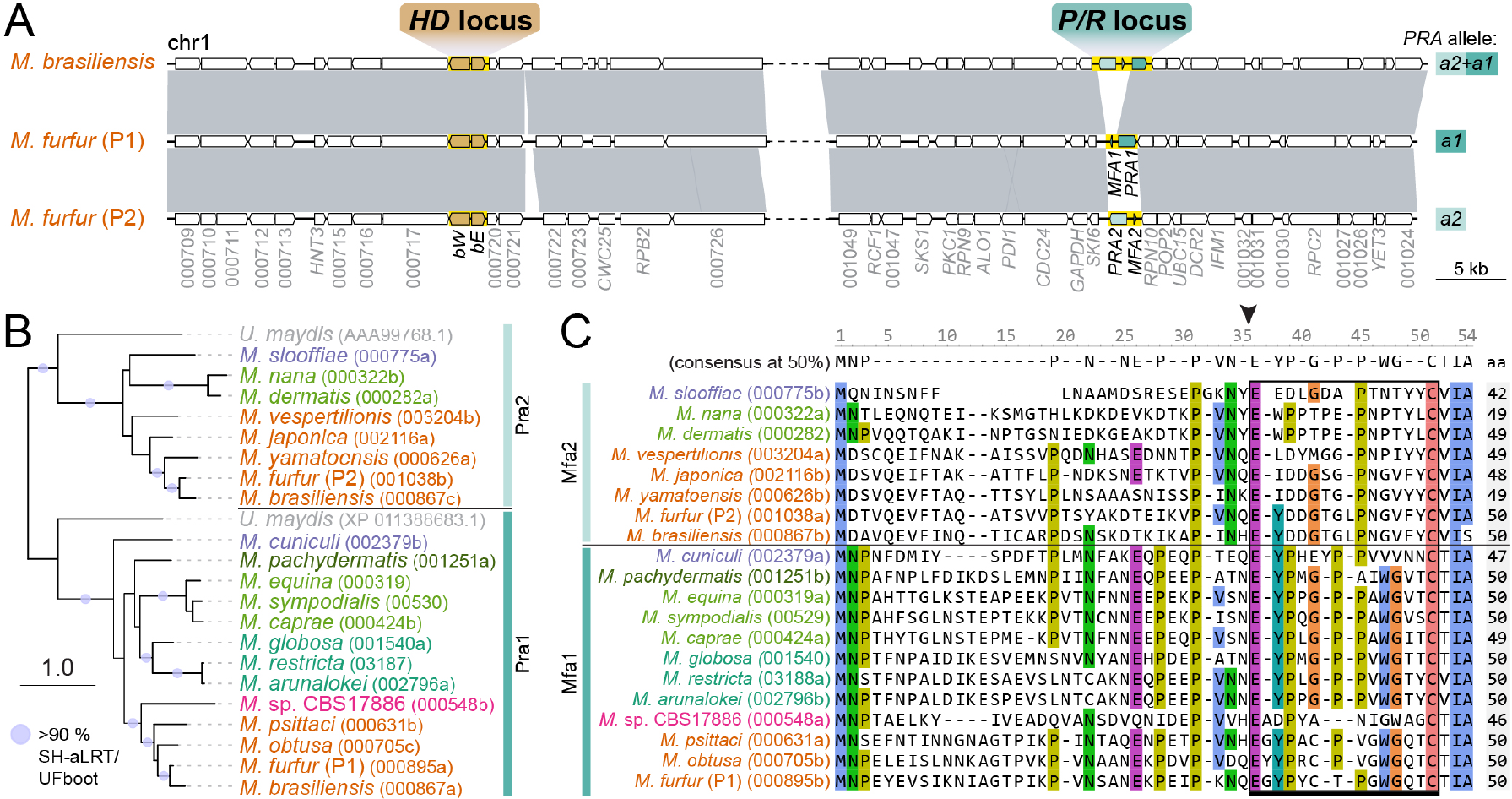
*MAT* loci structure and mating-type genes in *Malassezia*. (A) Synteny analysis comparing the chromosomal regions encompassing the *HD* and *P/R MAT* loci in two *M. furfur* representatives with different mating types and *M. brasiliensis*. The *P/R* and *HD* loci are highlighted in yellow. With the exception of *M. brasiliensis* that contains two alternate pheromone receptor genes (*PRA1* and *PRA2*) and one pheromone gene (*MFA2*) at the *P/R* locus, the gene structure of the two *MAT* loci is conserved across the genus (additional comparisons are in ***SI Appendix,* Fig. S4**). (B) Maximum likelihood phylogeny of *Malassezia* and *Ustilago maydis* Pra1 and Pra2 pheromone receptors showing clustering into two main groups representing the two alternate alleles. The tree was constructed with IQ-TREE2 (model LG+F+I+R6) and rooted at the midpoint. Internal branch support was assessed by 10,000 replicates of Shimodaira–Hasegawa approximate likelihood ratio test (SH-aLRT) and ultrafast bootstrap (UFboot) and well supported branches (>90%) are depicted by gray circles. Branch lengths are given in number of substitutions per site. (C) Sequence alignment of Mfa1 and Mfa2 pheromone precursors. The black arrowhead denotes the predicted cleavage site, giving rise to the peptide moiety of the mature pheromone (indicated by a black bar).

Next, synteny comparisons showed that both *MAT* loci are embedded within chromosomal regions that are largely conserved across species, although a few species-specific gene insertions/losses are observed in the neighboring regions, indicative of gene content remodeling as a byproduct of species-specific genome evolution (***SI Appendix*, Fig. S4**). A notable change in *M. brasiliensis* is that the *P/R* locus contains both alternate pheromone receptor genes (*PRA1* and *PRA2*), but only one pheromone gene (*MFA2*) (**Fig. 2A**). This suggests a scenario (illustrated in ***SI Appendix*, Fig. S5**) in which the *M. brasiliensis P/R* locus originated via intra-*MAT* recombination between opposite mating types. In summary, we conclude that despite the different chromosomal organization of *MAT* loci in *Malassezia* (pseudobipolar vs. tetrapolar), the gene content and structure is well conserved.

### Transitions in mating-type loci chromosomal organization in *Malassezia*

Examination of the species with a pseudobipolar configuration revealed a large range of distances between the two *MAT* loci, from an average of ∼145 kb in species of subclade B2, to ∼588 kb in species of clade A, and up to 871 kb in *M. slooffiae* (**Fig. 1D**). Along with the presence of tetrapolar species, this led to the initial hypothesis that the extant pseudobipolar configurations may have resulted from multiple independent events combining the two *MAT* loci on the same chromosome, as previously reported for tetrapolar-to-bipolar transitions in several basidiomycetes (23). Given the *Malassezia* species phylogeny (**Fig. 1A**), if we assume a tetrapolar ancestor and unidirectional transitions, we postulate five independent shifts from tetrapolar to pseudobipolar would have occurred during evolution (Model A; ***SI Appendix*, Fig. S6A**). Alternatively, the observed distance variation between the two *MAT* loci could imply departing from an ancestral pseudobipolar configuration followed by species- or clade-specific intrachromosomal rearrangements (such as large inversions) relocating the two loci within the chromosome. This latter scenario would imply that the tetrapolar *Malassezia* species are a derived state, in which case six independent transitions from pseudobipolar to tetrapolar would have had to occur to explain the extant distribution (Model B; ***SI Appendix*, Fig. S6B**). Lastly, we also considered the possibility of independent transitions occurring in both directions (Model C; ***SI Appendix*, Fig. S6C**).

The degree of gene synteny between extant species can reveal essential information about their evolution since their last common ancestor. To ascertain if one of the *MAT* locus evolutionary scenarios is more likely than the others, synteny blocks were reconstructed between all *Malassezia* genomes with SynChro (34), using *M. sympodialis* as reference (***SI Appendix*, Fig. S7**). We focused on comparisons of chromosomes containing *MAT* loci. This analysis revealed pseudobipolar species of clades B, C, and D share conserved synteny for most of the *MAT* chromosomes (**Fig. 3**). In *M. slooffiae* the greater distance between the two *MAT* loci is explained by two large inversions: one relocating the *HD* locus closer to the end of chr. 2 and the other inverting the relative positions of the centromere and the *P/R* locus (**Fig. 3B**). Hence, the similarities between the *MAT* chromosomes of three independently evolving lineages suggest a pseudobipolar *Malassezia* ancestor containing a *MAT* chromosome (henceforth referred as Anc1) structurally comparable to that of *M. sympodialis*.

**Fig. 3.**
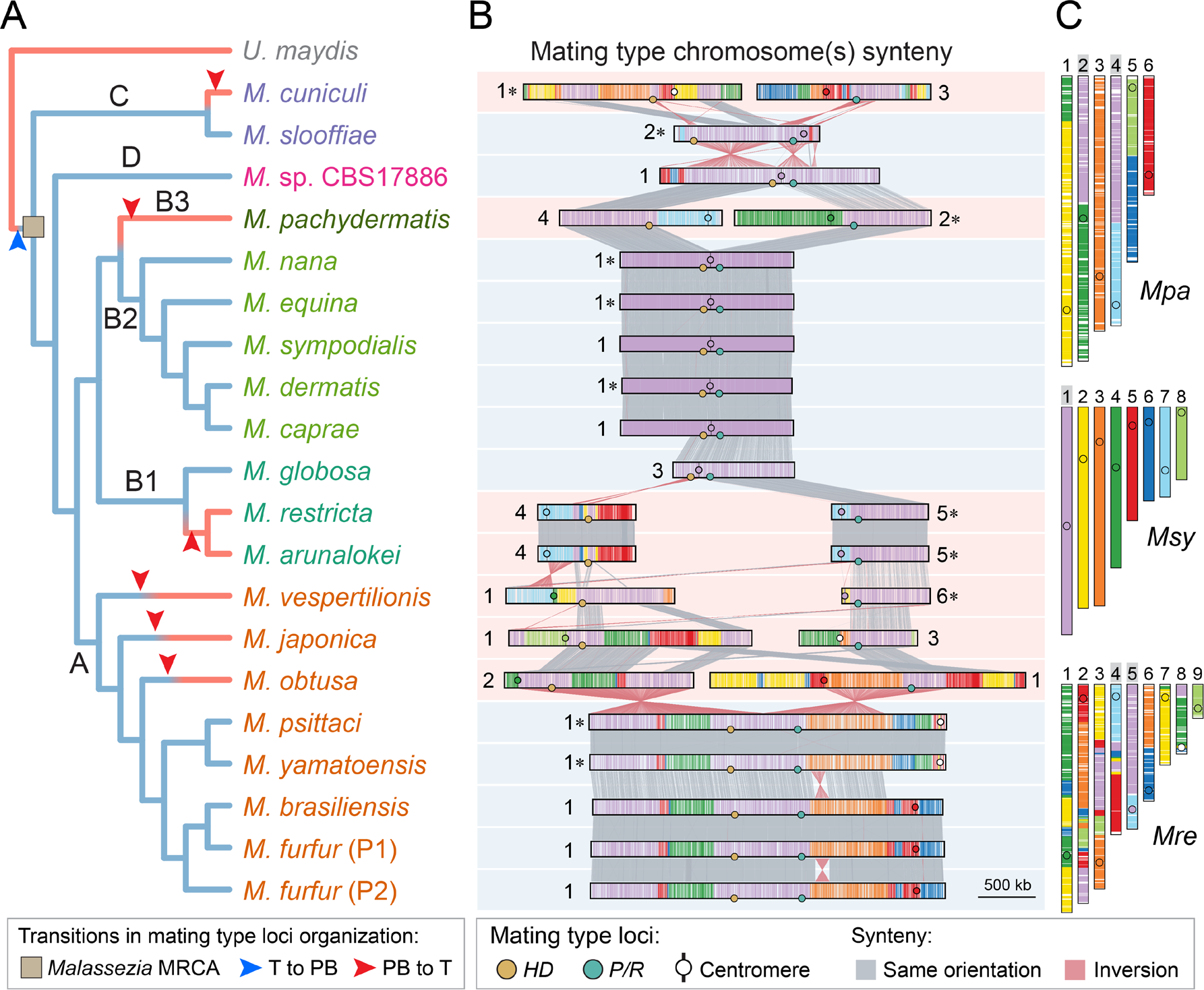
Proposed transitions in mating-type loci chromosomal organization in *Malassezia*. (A) Species-tree topology depicting hypothesized transition events in mating-type loci organization. A transition from tetrapolar (T) to pseudobipolar (PB) depicted by a blue arrowhead is proposed to have occurred only once in the *Malassezia* most recent common ancestor (MRCA) from which all the tetrapolar species have independently derived (depicted by red arrowheads in the tree branches). (B) Synteny analysis comparing the chromosomes containing the *HD* and *P/R MAT* loci across species. Chromosomes displayed in reversed orientation are marked by asterisks. (C) Chromosome structure differences between *M. sympodialis* (*Msy*; used as reference) and two other *Malassezia* species (*M. restricta* – *Mre*, and *M. pachydermatis* – *Mpa*) are shown as an example (refer to ***SI Appendix*, Figs. S6 and S7** for additional comparisons and other hypothesized models).

### Rearrangements associated with centromere breakage, or translocations in centromere-flanking regions, drove pseudobipolar to tetrapolar transitions

To investigate how pseudobipolar-tetrapolar shifts occurred, we compared *MAT*-containing chromosomes of the tetrapolar species within clades B, C and D with *M. sympodialis* chromosomes. On the basis of centromere predictions and synteny analysis, we hypothesize the transition in *M. pachydermatis* was triggered by a break at the centromere of the predicted Anc1 (**Fig. 4A**, and ***SI Appendix*, Fig. S8**). One of the acentric chromosome arms was rescued by fusion to an ancestral chr. 7 (Anc7), which is mostly structurally intact in *M. slooffiae*, *Malassezia* sp. CBS17886, all species of clades B2, and in *Malassezia japonica*/*M. obtusa* belonging to clade A. The other arm fused to an ancestral chr. 4 (Anc4) conserved in all species of clade B2, but more rearranged in others (**Fig. 4A** and ***SI Appendix*, Figs. S7** and **S8**). Besides relocating the two *MAT* loci onto different chromosomes, this event is associated with karyotype reduction from a predicted ancestor of clades B2-B3 with 8 chromosomes (16). An additional independent event giving rise to the extant 6-chromosome state in *M. pachydermatis* entailed telomere-telomere fusion of chrs. 6 and 8 conserved in clade B2 to generate chr. 5 of *M. pachydermatis* (**Fig. 3C** and ***SI Appendix*, Fig. S8C**). This event was apparently followed by a large inversion whose breakpoints map to the centromere- and telomere-proximal regions of *M. sympodialis* chr. 6. Supporting this series of events is the presence of two genes near the end of chr. 8 in *M. sympodialis*, which are found internalized in *M. pachydermatis* (***SI Appendix*, Fig. S8D**). Although not as yet definitively established, we posit this inversion may have inactivated *CEN6*, thereby stabilizing a presumably dicentric chromosome initially generated upon fusion.

**Fig. 4.**
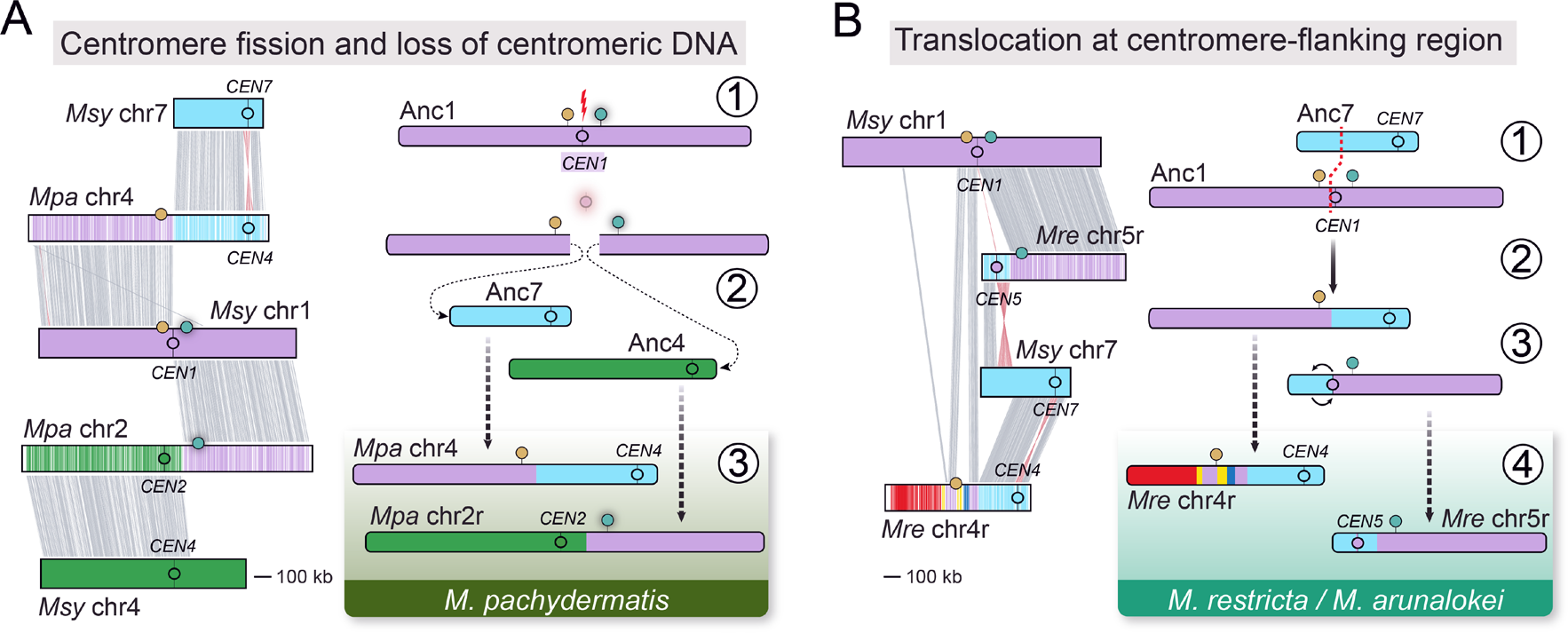
Main mechanisms underlying the pseudobipolar to tetrapolar transitions. (A) Synteny analysis (left side) between *M. sympodialis* and *M. pachydermatis MAT*-containing chromosomes indicates that the pseudobipolar to tetrapolar transition in *M. pachydermatis* possibly occurred due to centromere breakage on an ancestral chromosome (Anc1) containing the two *MAT* loci regions (*P/R*, green circle; *HD*, golden circle) (step 1). This event resulted in centromere loss, and the resulting chromosome arms were rescued by fusion to other chromosomes (step 2) thereby relocating the *P/R* and *HD* loci onto different chromosomes (step 3). (B) Differently, the transition in the common ancestor of *M. restricta* and *M. arunalokei* is hypothesized to have involved a translocation at the pericentromeric region of the Anc1 with a non-centromeric region of another chromosome (depicted here as the Anc7) (steps 1 and 2). Synteny analysis between *M. sympodialis* and *M. restricta* (left side) further indicate that an additional pericentric inversion occurred in the derived *P/R*-containing chromosome and further rearrangements reshaped the *HD*- containing chromosome during evolution (steps 3 and 4).

A second independent transition to tetrapolar occurred in clade B1, specifically in the common ancestor of the sibling species *M. restricta*/*M. arunalokei* (**Fig. 3A**). According to synteny analysis, this transition was initiated by a translocation between the centromere-flanking region of Anc1 and a non-centromeric region of Anc7, resulting in the progenitors of chrs. 4 and 5 of *M. restricta*/*M. arunalokei* containing, respectively, the *HD* and the *P/R* locus (**Fig. 4B**). A large pericentric inversion subsequently relocated *CEN5* while chr. 4 underwent further rearrangements (**Fig. 4B** and ***SI Appendix*, Fig. S9**).

Lastly, *Malassezia cuniculi* (tetrapolar) and *M. slooffiae* (pseudobipolar) are sister species but have diverged considerably since their last common ancestor. Their karyotypes differ markedly: *M. slooffiae* has nine chromosomes, representing the ancestral state of all *Malassezia* species (16), whereas *M. cuniculi* has only six (**Fig. 1A**). The *M. cuniculi* genome is also more rearranged (***SI Appendix*, Fig. S7**); this is particularly evident when comparing the *MAT*-chromosome of *M. sympodialis*, which is dispersed over the six chromosomes in *M. cuniculi* as opposed to only three in *M. slooffiae* (***SI Appendix*, Fig. S10**). Despite several rearrangements, many involving the surrounding centromeric regions of Anc1, the centromere itself remained intact, corresponding to *M. cuniculi CEN6* (***SI Appendix*, Fig. S10B**). Hence, the transition to physically unlinked *MAT* loci in *M. cuniculi* appears to be associated with a complex process of karyotypic reduction.

### Pseudobipolar structure in clade A is not secondarily derived

Although our analyses strongly suggest that the pseudobipolar configuration emerged in the *Malassezia* common ancestor, the presence of several tetrapolar species within clade A raised the possibility that a tetrapolar arrangement could be ancestral to this clade, and the pseudobipolar structure in the sibling species *M. yamatoensis/M. psittaci* and *M. furfur/M. brasiliensis* could have represented a more recent reversal to pseudobipolar (Model C in ***SI Appendix*, Fig. S6**). To investigate this, we first performed synteny analyses to model events leading to tetrapolarity of *M. vespertilionis*, which branched first within clade A (***SI Appendix*, Fig. S11**). We found a scenario remarkably similar to that described above for *M. restricta*/*M. arunalokei*, entailing a translocation in a centromere-flanking region. However, the translocation involved a different chromosome (chr. 2 of *M. sympodialis*) and the predicted translocation breakpoint localizes in the intergenic segment ∼5.7 kb upstream of the Anc1 centromere. The resulting chromosome harboring the *P/R* locus later underwent a pericentric inversion followed by breakage close to the centromere, which then healed by *de novo* telomere addition to generate an acrocentric chromosome (chr. 6 of *M. vespertilionis*; ***SI Appendix*, Fig. S11**). The tetrapolar state in *M. vespertilionis* therefore represents another independent transition from pseudobipolar to tetrapolar.

As for the other two tetrapolar species in clade A, *M. obtusa* and *M. japonica*, we first noted that their *P/R*- and *HD*-containing chromosomes share extensive blocks of synteny with the *MAT*- chromosome of pseudobipolar species within the *M. furfur*/*M. yamatoensis* clades (**Fig. 3B** and ***SI Appendix*, Fig. S12A**). Similarly, the *MAT*-chromosomes of *M. furfur* and *M. sympodialis*, have large blocks of conserved synteny, but the Anc1 centromere located between the two *MAT* loci was evicted from the *M. furfur*/*M. yamatoensis MAT*-chromosome (**Fig. 3B** and ***SI Appendix*, Fig. S12B**). Based on our analysis, this transition is best explained by an inversion and a translocation on Anc1 occurring after the split from the common ancestor with *M. vespertilionis*, but before the divergence of the other species (***SI Appendix*, Fig. S12C**). This inversion repositioned the *P/R* locus farther from the *HD* locus and moved the centromere closer to the end of the chromosome. The centromere was then exchanged via translocation between an ancestor of chr. 3 of *M. sympodialis* (Anc3) and the centromere-flanking region of the inversion-derived chromosome. One of the resulting products comprised most of Anc1 (including the two *MAT* loci) and a significant portion of Anc3, and likely represents the ancestral *MAT*-chromosome arrangement that emerged after divergence from the last common ancestor with *M. vespertilionis* (designated as AncA in ***SI Appendix*, Fig. S12C**). The other chromosome product inherited the centromere of Anc1 and is structurally identical to the extant chr. 4 of *M. furfur* (or chr. 5 in the *M. yamatoensis* clade) indicating it was retained in these pseudobipolar species. In *M. japonica* and *M. obtusa*, this second product underwent further rearrangements but retained the Anc1 centromere and neighboring regions (***SI Appendix*, Fig. S12D**). Finally, the proposed derived *MAT*-chromosome (AncA) also went through various species-specific rearrangements. One such rearrangement independently occurred in *M. japonica* and *M. obtusa*, and seemingly relocated the *P/R* and *HD* loci onto different chromosomes (tetrapolar transition), whereas in *M. furfur*/*M. yamatoensis* the two *MAT* loci were retained on the same chromosome (pseudobipolar) (***SI Appendix*, Fig. S12C**). Overall, this indicates that the pseudobipolar configuration is ancestral in *Malassezia*, and all extant tetrapolar species were independently derived from the pseudobipolar ancestral state.

### Engineered *Malassezia furfur* strains harboring compatible *MAT* alleles undergo hyphal development

Studies have demonstrated that *Malassezia* species harbor the key genetic components necessary for mating and meiosis, yet their sexual cycle remains elusive (5, 17, 18). In an effort to reevaluate the capacity of *Malassezia* for sexual reproduction, we first attempted inter-strain crosses within two *Malassezia* species, *M. furfur* and *M. sympodialis*, with incubation on various culture media. Specifically, fusion assays were initially carried out by mixing *M. furfur* strains CBS14141 *ade2*Δ::*NAT* (*MAT a2b1*) x CBS9369 (*MAT a1b2*) (***SI Appendix*, Table S2**). The co-cultures were resuspended in water and plated onto modified minimal medium (mMM, see ***SI appendix***, **SI Material and Methods** for details) supplemented with nourseothricin to select for adenine prototrophy and NAT resistance, respectively. However, despite prolonged incubation, no colonies were obtained. Similarly, *M. sympodialis* strains ATCC42132 NAT^R^ (*MAT a1b1*) x ATCC44340 NEO^R^ (*MAT a2b2*) (***SI Appendix*, Table S2**) were mixed on various culture media and inoculated onto mMM media supplemented with nourseothricin and neomycin/G418, but also in this case no fusion products or meiotic recombinants were recovered.

Next, a larger collection of genetically characterized *M. furfur* strains (26) (***SI Appendix*, Table S2** and **Fig. S13A**) were crossed in pairwise combinations and examined by microscopy for the possible presence of sexual structures typical of basidiomycete fungi, such as hyphal formation, clamp connections, teliospores, basidia, and basidiospores. Despite our efforts, we were unable to observe any signs of mating in these cases. To circumvent this, we took a molecular approach and generated a genetic transgene comprising the two genes of the *P/R a1* locus (*PRA1* and *MFA1*) derived from strain CBS9369, a *NEO* selectable marker, and the two genes of the *HD b4* locus (*bW4* and *bE4*) obtained from strain CBS7710. This transgene was then introduced via *Agrobacterium tumefaciens*-mediated transformation (35) into the presumably compatible *M. furfur* strain CBS14141 (*MAT a2b1*) (**Figs. 5A** and **5B**, and ***SI Appendix*, Fig. S13C**). One NEO^R^ transformant confirmed by PCR to contain both native and exogenous *MAT a* and *b* alleles (***SI Appendix*, Figs. S13D-E**) was chosen for further analysis. Remarkably, this *M. furfur* transformant produced hyphae on various agar media (stronger on YPD and YT, and fewer hyphae on MS, FA, PDA, and V8 pH 7), whereas no hyphal growth was observed in the wild-type (WT) strain CBS1414 under these conditions (***SI Appendix*, Fig. S14A**). This *M. furfur* transformant (*MAT a1a2b1b4*) was named *A*on *B*on (or “solo”; **Fig. 5B**), following the nomenclature used for the *U. maydis* solopathogenic strain SG200 engineered to undergo hyphal development and cause smut disease in maize without a mating partner (36). Additionally, when grown in both solid or liquid YPD supplemented with compounds reported to promote hyphal formation in *Malassezia* (37) (squalene, potassium nitrate, sodium chloride, ferrous sulphate, magnesium sulphate, and glycine; hereafter referred as “YPD + supplements”; see ***SI Appendix***, **SI Material and Methods**), the *M. furfur* “solo” transformant displayed robust hyphal growth featuring branches and septa, and both fused and unfused clamp-like structures (**Fig. 5E** and ***SI Appendix*, Fig. S14B-C**). Conversely, the *M. furfur* WT produced only yeast cells on the same growth media (**Fig. 5D** and ***SI Appendix*, Fig. S14B**). In *U. maydis*, similar clamp-like structures are formed during the dikaryotic stage and participate in nuclear migration (38). Although the addition of these supplements enhanced the formation of hyphae and clamp-like cells, these structures were also observed, albeit less frequently, in liquid YPD media without the addition of supplements. Last, we noticed that the addition of lipids (e.g., tween and ox-bile) did not affect the ability of the *M. furfur* “solo” transformant to produce hyphae.

**Fig. 5.**
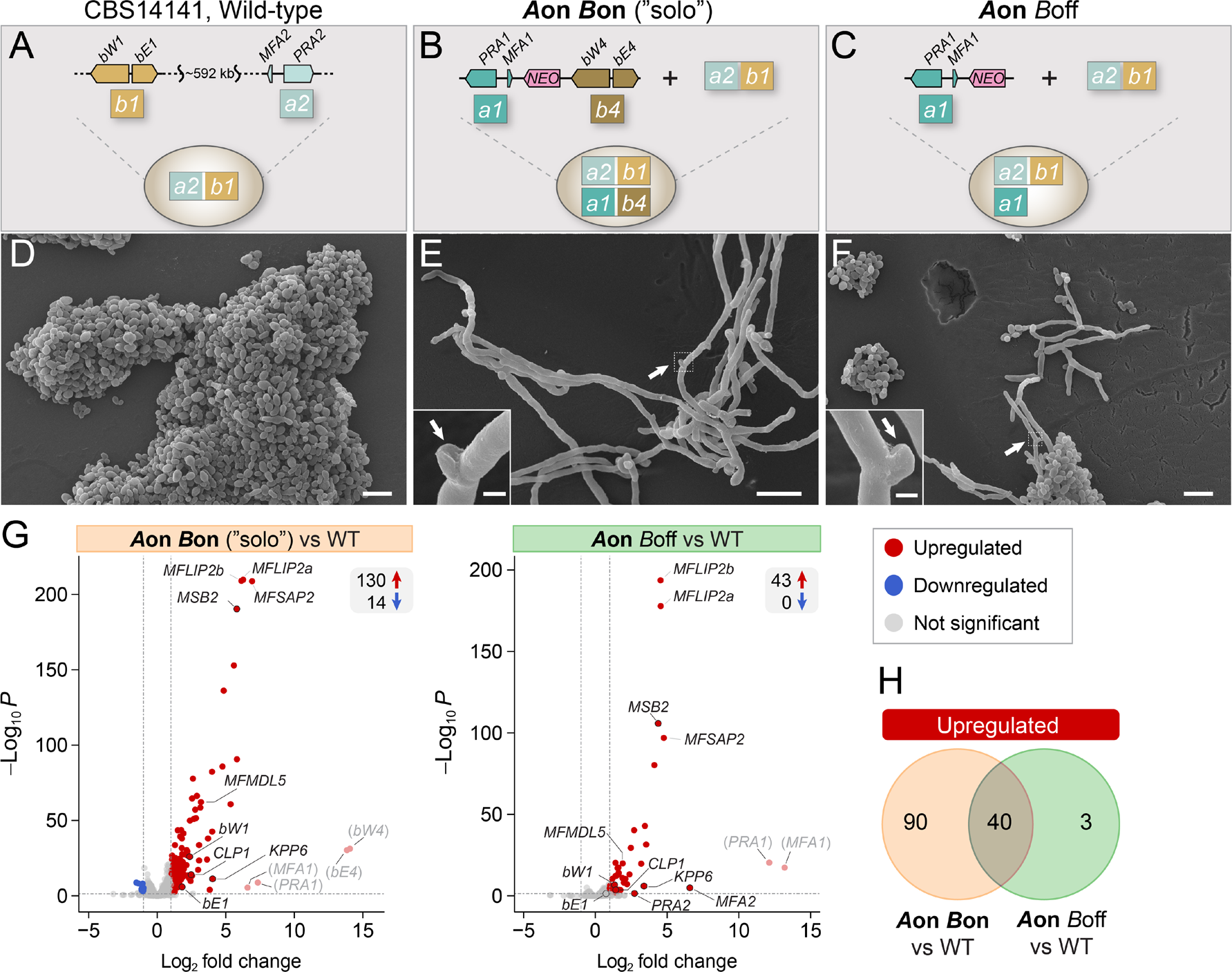
*Malassezia furfur* strains expressing compatible *P/R* and *HD* genes are self-filamentous and resemble the initial stages of sexual development of other basidiomycetes. (A-C) The genetic constructs *a1*-*NEO*-*b4*, and *a1*-*NEO* were introduced into the recipient *M. furfur* wild-type (WT) haploid strain CBS14141 (*MAT a2b1*) to generate the *A*on *B*on “solo” (*MAT a1a2b1b4*) and *A*on *B*off (*MAT a1a2b1*) strains, respectively. (D-F) Scanning electron microscopy images of *M. furfur* WT only growing as yeast, and transformants expressing a compatible *P/R* and *HD* allele (*A*on *B*on, “solo”) or a compatible *P/R* allele (*A*on *B*off) showing hyphal development (self-filamentation) and the formation of clamp-like cells (indicated by white arrows). Scale bars = 10μm (1μm in the insets). (G) Volcano plot representations of differentially expressed genes. Red and blue points represent, respectively, transcripts with significantly increased or decreased levels in *M. furfur* transformants relative to the WT, for Log2 fold change (FC) ≥ ±1, and a false discovery rate (FDR) < 0.05. Grey dots represent transcripts with no significant changes. Faded red dots representing the expression levels of *PRA1*, *MFA1*, *bW4* and *bE4* are depicted only as internal controls as their upregulation is expected given their absence in the parental strain (i.e., only harbors the allele pairs *PRA2*/*MFA2* and *bW1*/*bE1*) Outlined genes are predicted to have a role in sexual reproduction. (H) Venn diagram showing common and unique sets of upregulated genes between the comparisons reported in panel G.

To dissect the contribution of each *MAT* locus (*P/R* and *HD*) in the yeast-to-hyphae transition, we then sought to individually introduce the *MAT a1* or *b4* genes into *M. furfur* CBS14141 WT strain (***SI Appendix*, Fig. S14C)**. Transformants carrying the *MAT a1* genes were readily obtained, and PCR analysis confirmed the presence of both the native *MAT a2* locus and the exogenously introduced *MAT a1* (***SI Appendix*, Fig. S13D-E**). A *MAT a1a2b1* strain carrying two compatible alleles of the *P/R* locus was selected and named *A*on *B*off (**Fig. 5C**). Curiously, all transformants generated by introducing the *MAT b4* genes alone were unable to maintain stably G418/neomycin genetic resistance and were not analyzed further. When grown on YPD agar + supplements the *M. furfur A*on *B*off strain displayed characteristics analogous to the “solo” strain, including hyphae with septa and clamp-like structures (**Fig. 5F** and ***SI Appendix*, Fig. S14**). Therefore, no obvious morphological differences were observed between the *A*on *B*off and the “solo” mutant under the conditions tested. Taken together, the hyphal growth observed in these *MAT* engineered *Malassezia* isolates seems to represent the initial stages of sexual development and this genetic analysis thus provides evidence for a role of the mating-type genes in controlling this process.

### Key mating genes are upregulated in *M. furfur* “solo” and *A*on *B*off strains exhibiting hyphal growth

To assess if the hyphal phenotype observed in *M. furfur* “solo” and *A*on *B*off transformants is associated with activation of genes known to function during sexual reproduction in other basidiomycetes, the transcriptomic profiles of these strains were compared to the *M. furfur* CBS14141 WT, all grown under conditions that promoted hyphae formation (liquid YPD + supplements). RNA-Seq analysis using as threshold a false discovery rate (FDR) lower than 5% and log2 fold change (FC) ≥ ±1.0 identified 43 and 144 differentially expressed genes (DEGs) in the *A*on *B*off and “solo” strains compared to the WT, respectively, from 4,601 predicted genes (**Fig. 5B**, ***SI Appendix*, Fig. S15**, and **Dataset S2**). Among the upregulated DEGs, three are found solely in the *A*on *B*off strain, 90 genes are exclusive to the “solo” strain, and 40 genes are shared by both *A*on *B*off and “solo” strains (**Fig. 5C**). Although a gene ontology enrichment analysis did not reveal any significant enrichment among the DEGs in both the *Aon B*off and “solo” strains, several noteworthy genes were identified and are discussed in the following sections.

In the *A*on *B*off strain, three genes were specifically upregulated: GLX27_002835, encoding an unknown *Malassezia*-specific protein, and *MFA2* (GLX27_001038a; log2 FC = 6.59) and *PRA2* (GLX27_001038b; log2 FC = 2.73), encoding the endogenous mating-type pheromone and receptor respectively, both expressed at very low levels in the WT strain (**Dataset S3**). The observed upregulation of *MFA2* and *PRA2* in the *A*on *B*off strain suggests that a compatible interaction was established with the products of the introduced and expressed pheromone (*MFA1*) and receptor (*PRA1*) genes (**Datasets S3** and **S4**). In contrast, the “solo” strain shows no induction for *MFA2* and *PRA2*, with these genes remaining at WT expression levels. In contrast to the *A*on *B*off strain, the “solo” strain additionally expressed the *bW4* and *bE4* genes (*b4* allele) introduced by transformation, and the native *bW1* and *bE1* genes (*b1* allele) were both upregulated in this context (**Fig. 5B** and **Dataset S2**). This presumably allows the formation of an active bW/bE heterodimer, which is likely to negatively affect the transcription levels of the pheromone and receptor genes, as reported in *U. maydis* and *C. neoformans* (39–41).

Among the 40 genes shared and upregulated in both the *A*on *B*off and “solo” strains, *Malassezia* orthologs of three significant genes in *U. maydis* sexual or pathogenic development were identified: *CLP1* (GLX27_003783), *KPP6* (GLX27_002769), and *MSB2* (GLX27_000860).

*CLP1* (<u>cl</u>am<u>p</u>less <u>1)</u> is required for clamp-like cell formation and initiation of *U. maydis* pathogenic development (38). *KPP6* encodes a MAP kinase that is active post cell-fusion in dikaryotic hyphae and is upregulated by the bE/bW heterodimer. *U. maydis kpp6* mutants develop appressoria (a structure required for plant penetration) but cannot invade the host plant (42). Similarly, the product of *MSB2* is a transmembrane mucin that senses plant surface cues and is essential for appressoria formation in *U. maydis*, acting upstream of the MAP kinase signaling pathway (43). Hence, the enhanced expression of these genes in the *M. furfur A*on *B*off and “solo” strains relative to WT suggests that they may fulfill key roles in the initiation of the sexual cycle and hyphal development in *Malassezia*.

In addition, four genes were upregulated in both *A*on *B*off and “solo” strains, which could be important for *Malassezia* physiology and pathogenicity. The first set comprises three highly upregulated genes encoding secreted lipases (**Fig. 5B** and **Dataset S2**). Two of these genes, *MFLIP2a* and *MFLIP2b*, are adjacent and result from a gene duplication within clade A (***SI Appendix*, Fig. S16**). The proteins they encode contain a secretory lipase domain belonging to the LIP family (PF03583). The third gene, *MFMDL5*, encodes a protein with a lipase class 3 domain (PF01764; ***SI Appendix*, Fig. S17**) and its counterpart in *M. globosa* (Mgmdl5) can utilize mono-, di-, and tri-glyceride substrates (44). Lastly, we observed a striking upregulation of *MFSAP2* (GLX27_000215; log2 FC = 4.77 and 6.92, for *A*on *B*off and “solo”, respectively) encoding a secretory aspartyl protease. This gene is one of five secretory aspartyl protease (SAP) genes previously identified in *M. furfur* (45) and our own survey indicates that there may be up to 9 SAP-like genes in this species (***SI Appendix*, Fig. S18**). A recent study revealed that *MFSAP2* expression is essentially undetectable in *M. furfur* CBS14141 WT, whereas *MFSAP1* (GLX27_000216), which lies adjacent to *MFSAP2* in the genome, is the most highly expressed SAP gene (46). Our data confirmed that although *MFSAP1* is highly expressed, its expression does not differ between the mutant and WT strains (GLX27_000216; **Datasets S2** and S3). *M. furfur MFSAP1* seems to play an essential role in fungal skin colonization and tissue inflammation, and a compensatory expression of the other SAP genes was not observed when *MFSAP1* was deleted (46). Therefore, the increased expression of *MFSAP2* and three lipase genes in the *A*on *B*off and “solo” strains could represent a life cycle stage-specific response possibly associated with hyphal growth during sexual development.

## Discussion

Despite a surge in knowledge about *Malassezia* over the last two decades, the natural history, biology, and epidemiology related to colonization and infection by this group of fungi remains largely unexplored. One major gap in our knowledge pertains to the sexual cycle of *Malassezia* species, which despite mounting genomic evidence suggesting mating and meiosis capabilities, remains elusive. Our analyses revealed six independent instances of derived tetrapolarity, wherein the *P/R* and *HD* mating-type loci were secondarily relocated onto separate chromosomes from an ancestrally inferred pseudobipolar configuration. Furthermore, by introducing compatible *P/R* and *HD* alleles into *M. furfur*, we engineered a strain capable of initiating a developmental hallmark of basidiomycete sexual reproduction, thereby providing evidence for the presence of an extant sexual cycle in this remarkable group of yeasts.

Building on prior studies, the phylogenetic relationships within the *Malassezia* genus were more clearly resolved by integrating whole-genome data, and a new *Malassezia* species isolated from bats was uncovered. Leveraging this high-quality genomic dataset, two genomic features were explored: telomeres and mating-type loci chromosomal organization.

Our research revealed that telomeric sequences in *Malassezia* consist of tandem arrays of one of four identified telomere-repeat motif variants. Similar diversity in telomeric repeat sequences has been observed in related fungal species, particularly in ascomycetous yeasts, where sequence and length variability are even more pronounced and accompanied by rapid evolution of cognate telomere-binding proteins (47, 48). Future identification of the telomerase RNA component in *Malassezia* (a long noncoding RNA that serves as a short template for telomere replication by the telomerase reverse transcriptase catalytic component), similar to recent studies in *U. maydis* (49), could further illuminate these telomere sequence differences. Besides telomeric variability, some species contain inverted repeats after the telomeric repeat tract, predicted to encode non-LTR retrotransposons that might insert into telomeres. While the function and activity of these candidate retrotransposons is presently unknown, studies in other fungi and beyond indicate these elements may cause chromosome instability, yet also provide adaptive benefits (29, 32, 50). In *Drosophila*, the loss of the telomerase gene is associated with the replacement of telomeric sequences by tandem retrotransposons, which suggests these elements may prevent chromosome shortening when telomerase function is compromised (51). These findings reveal plasticity of *Malassezia* chromosome ends, hinting at a contribution of these regions in shaping the *Malassezia* genomes, with potential effects on speciation and adaptation.

The ancestor of all basidiomycetes likely had unlinked *P/R* and *HD* mating-type loci (tetrapolar), each composed of alternate alleles of a single pair of genes controlling pre- and post-mating mating compatibility (23). However, multiple basidiomycete genera (*Ustilago*, *Cryptococcus*, *Trichosporon*, *Microbotryum*) demonstrate *P/R*-*HD MAT* loci linkage (bipolar) formed through chromosomal translocations or fusions of the ancestral *P/R* and *HD* chromosomes (52–58). This derived state is usually associated with the formation of a large nonrecombining region encompassing multiple genes that are co-inherited during meiosis (24). A recent model proposes that these nonrecombining regions can expand in a stepwise manner due to sheltering of recessive alleles by inversions linked to a permanently heterozygous locus (such as *MAT*), and become highly rearranged between mating types (59). This is reinforced in the case of highly selfing/inbreeding mating systems, where mating occurs preferentially among meiotic products from a single diploid individual. However, the extent to which recombination suppression occurs beyond *MAT* is hypothesized to depend on the duration of the haploid phase (as it enhances purging of deleterious recessive mutations) and the rate of outcrossing (59).

Here we demonstrate that the genetic makeup of the *P/R* and *HD* mating-type loci is conserved across *Malassezia* species, analogous to the archetypal basidiomycete *MAT* loci, and consistent with heterothallic mating. Importantly, by reconstructing synteny blocks between all *Malassezia* genomes, along with centromere identification, we inferred that the *Malassezia* common ancestor was most likely composed of two *MAT* loci located some distance apart on the same chromosome (pseudobipolar) with a centromere located between the two regions. It is uncertain how this ancestral arrangement was established, but this likely involved chromosomal rearrangements, such as translocations, as proposed for other tetrapolar-to-bipolar transitions (52–58). However, this ancestral pseudobipolar arrangement, still seen in most *Malassezia* species albeit with a few modifications, deviates from the typical bipolar configuration. Indeed, at least in *M. sympodialis* (17, 18) and *M. furfur*, for which genome sequences of additional isolates are available, synteny is completely conserved along the *MAT* chromosome between opposite mating types. Consistent with these observations, past studies found alternative allelic combinations of the *P/R* and *HD* loci among isolates of the two species, implying that recombination between the two *MAT* loci occurs (17). In this model, we hypothesize that chromosomal rearrangements within the *P/R*-*HD* intervening region (e.g., fissions or translocations) are not as detrimental as they would be under a typically bipolar configuration with linked *P/R* and *HD* loci. In agreement, we identified six independent transitions from pseudobipolar to tetrapolar, with five cases involving translocations near the centromere of the ancestral *MAT* chromosome, and one case (*M. pachydermatis*) involving centromere breakage followed by fusion of the acentric chromosome arms to other chromosomes. Centromere breakage is a known mechanism of karyotypic change in *Malassezia* (16), and similar survival by fusion of acentric chromosome arms after centromere fission has been observed in *Schizosaccharomyces pombe* (60) as well as in *Cryptococcus deuterogattii* upon centromere deletion (61). Importantly, and in contrast to other basidiomycetes that usually have large regional centromeres (> 15 kb, often repeat rich), *Malassezia* species have short (< 5 kb-long) regional centromeres, and by virtue of their AT-richness, they seem to constitute fragile sites in the genome (16). We therefore posit that centromeres and, possibly their nearby flanking regions, are more prone to undergo breakage and interchromosomal rearrangements and are thus involved in the repeated transitions from pseudobipolar to tetrapolar in *Malassezia*.

The repeated transitions to linked *MAT* loci in basidiomycetes are suggestive of strong selection likely favoring increased gamete compatibility in species with a selfing/inbreeding mating system. Indeed, as mating can only occur when alleles differ at both *MAT* loci, gametes are compatible with only 25% of other gametes in species with two *MAT* loci and two alleles each (tetrapolar), while these odds rise to 50% in species with a single *MAT* locus and two alleles (bipolar). In comparison, the pseudobipolar configuration observed in *Malassezia* results in compatibility odds that are likely between those observed for the bipolar and tetrapolar configurations and depend on the frequency of a single cross-over occurring between the two *MAT* loci during meiosis. Thus, the pseudobipolar system does not seem to be under the same types of selection and constraints as observed in other bipolar basidiomycetes with linked *P/R* and *HD* loci, potentially delaying a transition to a strictly bipolar state. The lack of recombination within bipolar *MAT* alleles of some basidiomycetes (e.g., in *C. neoformans*, the barley smut *U. hordei*, and anther-smut *Microbotryum* species) has also facilitated accumulation of transposable elements in these regions due to a reduced efficacy of selection against deleterious insertions. However, *Malassezia* genomes have a very low density of transposable elements (5), which may pose a further hindrance to formation/stabilization of non-recombining regions. Future studies comparing sequence diversity of *MAT* and adjacent regions across multiple *Malassezia* species and strains, will be key to further understanding the evolution of these chromosomal regions.

The ability to switch between yeast and hyphal growth is well recognized in numerous fungal pathogens of plants and animals including *U. maydis*, *C. neoformans*, and *Candida albicans* and is known to be associated with virulence in some of these species (22, 62, 63). “Shapeshifting” between different morphologies in human fungal pathogens may provide advantages for nutrient acquisition, spread into new areas of the host, and evade the immune system (62). Dimorphism in *Malassezia* was first noticed based on observation of both yeasts and short hyphal filaments on lesional skin sites of patients with pityriasis versicolor (64) (described as the “spaghetti” and “meatballs” appearance). Since then, the presence of *Malassezia* hyphal forms has been reported for several species in both laboratory and clinical settings, with a recent report suggesting that they might be a factor contributing to pathogenesis of seborrheic dermatitis (65). This is intriguing because the hyphal cell type also coincides with the sexual development in dimorphic basidiomycete fungi, which is usually initiated upon mating of compatible partners and regulated by the mating-type genes (23). However, none of the aforementioned studies in *Malassezia* examined the relationship between the *MAT* gene composition and dimorphism.

Mating in *Malassezia* has never been achieved under laboratory conditions, although hybrid strains of *M. furfu*r have been isolated from nature and they are proposed to originate from mating events (26). Our inter-strain crossing attempts in both *M. furfur* and *M. sympodialis* under different conditions were not fruitful, potentially due to the media used not being conducive for mating, a requirement for mating on the host, or other unidentified genetic constraints. To overcome this, we engineered a haploid *M. furfur* strain harboring compatible *MAT* alleles (*A*on *B*off or *A*on *B*on “solo”). Our approach provides several lines of evidence for extant sexual development in *Malassezia* beyond the identification of mating and meiotic genes and demonstrates an association between yeast-to-hyphae transition and compatibility at the *MAT* loci. First, engineered haploid *M. furfur* strains grow as hyphae, unlike the haploid WT strain that grows only as yeast. Second, the branched hyphal filaments produced by these strains display clamp-like structures typical of a basidiomycete sexual stage. Third, mating-type determining genes are upregulated in *A*on *B*off and “solo” strains suggesting compatibility leading to activation of downstream pathways involved in pheromone response and morphogenesis in other fungi. In the “solo” strain we hypothesize bE/bW heterodimerization generates an active transcriptional regulator that activates/represses other genes, directly or indirectly through interaction with other proteins. Finally, upregulation of *CLP1* in both *A*on *B*off and “solo” strains may mediate steps in sexual reproduction as in in *U. maydis* and *C. neoformans* (38, 66).

Our transcriptomic analysis further suggests that the dimorphic transition in *M. furfur A*on *B*off and “solo” strains may contribute to skin colonization and pathogenicity. This is underscored by the upregulation of three secreted lipases, adding to the arsenal of lipases expressed and secreted by *M. furfur* yeast cells. Lipases secreted by *Malassezia* breakdown skin sebum-derived lipids (e.g., mono-, di-, and triglycerides) into saturated and unsaturated fatty acids. The saturated fatty acids are taken up by *Malassezia* for growth and proliferation, whereas the irritating unsaturated fatty acids accumulate on the outermost layer of the skin (stratum corneum). This accumulation, which we hypothesize could be further enhanced during hyphal growth, may in turn disrupt permeability of the skin barrier and lead to skin disorders (67). Additionally, both *M. furfur A*on *B*off and “solo” strains showed markedly increased expression of *MFSAP2*, which encodes one of nine putative secretory aspartyl proteases. Recent studies demonstrated that *MFSAP1*, which lies adjacent to *MFSAP2*, is the dominant secretory aspartyl protease in *M. furfur* WT and has critical roles in controlling cell adhesion during colonization and inducing inflammation in barrier-compromised skin (46). Therefore, the *Malassezia* hyphal growth detected associated with initiation of sexual development may have important roles in skin colonization and inflammation, especially in patients with impaired skin barriers, and thus affect its pathogenicity and virulence.

Collectively, our work reveals fundamental aspects of *Malassezia* biology. The genomic resources now available for *Malassezia* spp. can be leveraged to identify differences between species and strains, and the characterization of the *MAT* loci across species can assist in determining if different mating types co-exist and thrive on the same host individual. The role of specific genes in mating and hyphae formation can now be directly assessed, applying gene deletion methods recently developed for *Malassezia* (35). Analysis of the hyphae-producing strains in a murine model of *Malassezia* colonization (68) would be an essential next step to gain insight into the role of *Malassezia* cell morphology in skin colonization and disease, and investigate if the skin provides an environment specifically conducive to mating and sexual reproduction.

## Materials and Methods

Full details are given in ***SI Appendix***. Briefly, selected *Malassezia* strains were cultured in modified Dixon’s media or Leeming & Notman modified media at optimum growth temperature. High-molecular weight DNA for Nanopore sequencing was obtained by a customized cetyltrimethylammonium bromide (CTAB) extraction procedure (69). For Nanopore sequencing, barcoded libraries were obtained using unfragmented DNA and pooled together in MinION or PromethION R9.4.1 flow cells and sequenced. Basecalled reads passing filtering were de- multiplexed and assembled with Canu v2.1.1 (70). Accuracy of draft assemblies was improved with Medaka v1.5.0 or Nanopolish v0.11.2, and further polished with Pilon v1.22 (71) using Illumina paired end reads. The draft assemblies were validated by inspecting read coverage profiles with the Integrative Genomics Viewer after realigning the Canu-corrected and Illumina reads. Gene prediction and annotation on repeat-masked assemblies was performed with the funannotate pipeline v1.8.9, guided with well-annotated proteins of *M. sympodialis* and with RNA-seq where available.

*Malassezia* genomes were analyzed and compared via several approaches. For species-tree reconstruction, single copy orthologs were identified across *Malassezia* spp. and the outgroup *U. maydis* with OrthoFinder v2.5.4 (72) and aligned with MAFFT v7.310 (73). A maximum likelihood tree was constructed using a concatenation with gene-based partitioning approach in IQ-TREE v2.1.3 (74) with 1,000 ultrafast bootstraps replicates, and a gene-based coalescence tree was obtained with ASTRAL-MP v5.15.2 (75). Chromosome ends were inspected for the presence of telomeric and other repeats that co-occurred over all or most of the contigs. Centromeric regions were predicted by measuring GC-content in sliding windows of 250 bp (step size = 2) and identifying the corresponding GC-minimum in each contig. GC-content variation and distribution were evaluated with OcculterCut (76) with default settings. Conserved synteny blocks were identified between pairwise combinations of genomes with SynChro (34) setting the block stringency delta to three, and linear synteny plots comparing *MAT* loci and *MAT*-containing chromosomes were performed with Easyfig v2.2.3 (77).

*Malassezia furfur* strains harboring different combinations of *MAT* alleles (*MAT a1a2b1, A*on *B*off or *MAT a1a2b1b4*, *A*on *B*on “solo”) were obtained by *A. tumefaciens*-mediated transformation of *M. furfur* CBS14141 WT with plasmids containing transgenes with the *a1* or the *a1* and *b4* alleles together with a *NEO* marker, and assembled within the T-DNA region of plasmid pGI3 through *in vivo* recombination in *Saccharomyces cerevisiae* (35) (plasmids pGI26 and pGI52, respectively). The resulting transformants and WT were analyzed by light and scanning electron microscopy and subjected to RNA-Seq analysis. RNA extraction of *M. furfur* CBS1414 WT, *A*on *B*off, and “solo” transformants was performed with TRIzol as previously described (15) from three independent cultures grown at 30°C for 2 days in YPD + supplements. To compare the transcriptomic profiles of *A*on *B*off and “solo” transformants relative to WT, 50-bases single end Illumina reads were filtered and trimmed with Trim Galore v0.6.7 and mapped with STAR v.2.7.4a to a reference genome combining the *M. furfur* CBS14141 nuclear genome and the *a1*-*NEO*-*b4* transgene sequence. Read count matrices were obtained with featureCounts v2.0.1 and differential expression assessed with DESeq2 v1.36.0 (78) using a FDR < 0.05 and log2 FC > ± 1.0.

## Acknowledgments

We thank Drs. Minou Nowrousian, Salomé LeibundGut-Landmann, and John F. Rawls, for critical reading of the manuscript and insightful suggestions. We thank Anna Floyd Averette for technical support, and Dr. Fred Dietrich for access and assistance with computational resources. We also wish to thank Dr. Thomas L. Dawson for generously providing Illumina data for *M. psittaci* and *M. brasiliensis*. This study was supported by NIH/NIAID R01 Grants AI039115-26 and AI050113-18, awarded to J.H. and R21 Grant AI168672-01, awarded to J.H. and Salomé LeibundGut-Landmann. *Malassezia* work in the G.I. lab is supported grant LF-OC-22-001060 from the LEO foundation. K.S. acknowledges the JC Bose National Fellowship (Science and Engineering Research Board, Govt. of India, JCB/2020/000021). J.H. is Co-Director and Fellow of the CIFAR program Fungal Kingdom: Threats & Opportunities. Any use of trade, firm, or product names is for descriptive purposes only and does not imply endorsement by the U.S. Government.

