## Supporting Information for "Frequent transitions in mating-type locus chromosomal organization in *Malassezia* and early steps in sexual reproduction"

**This PDF file includes:**

Extended Materials and Methods
Figures S1 to S18
Tables S1 to S3
Legends for Datasets S1 to S4
Disclaimer
SI References

**Other supporting materials for this manuscript include the following:**

Datasets S1 to S4

**Extended Material and Methods**

**Strains and growth conditions.** *Malassezia* strains of different species (Tables S1 and S2) were routinely grown and maintained on either modified Dixon's (mDixon) medium [3.6% malt extract (Bacto), 1% mycological peptone (Oxoid), 1% ox bile (Oxoid), 1% v/v Tween 60 (Sigma), 0.4% glycerol, 2% Bacto agar] or in modified Leeming & Notman (mLNA) media (1% Bacteriological peptone, 1% glucose, 0.2% yeast extract, 0.8% ox bile, 1% glycerol, 0.05% glycerol monostearate, 0.5% v/v Tween 60, 2% Olive oil, 1.5% Bacto agar) at their optimum growth temperatures. *Saccharomyces cerevisiae* strain FY834 [*MAT*α; *his3*; *ura3*; *leu2*; *lys2*; *trp1* (1) was used to generate plasmids for *Agrobacterium tumefaciens*-mediated transformation (ATMT) through *in vivo* recombination, as previously described (2). Selection of *S. cerevisiae* transformants was carried out on yeast nitrogen base (YNB) minimal medium without uracil, supplemented with histidine, leucine, lysine, and tryptophan. *A. tumefaciens* strain EHA105 used for ATMT was maintained at 30˚C on Luria Bertani (LB).

**Genomic DNA extraction.** High-molecular weight genomic DNA was prepared using a customized cetyltrimethylammonium bromide (CTAB) extraction as previously described (3). Briefly, the cell pellet from a 100-mL culture grown as described above was recovered by centrifugation, frozen at -80˚C overnight and lyophilized. The pellet was then reduced to a fine powder by vortexing with approximately 5 mL of glass beads (3 mm), and 40 mL of CTAB lysis buffer was added, followed by a 1h incubation at 65°C. After cooling the mixture on ice, the glass beads were discarded, and the suspension was gently mixed by inversion with an equal volume of chloroform. After refrigerated centrifugation at maximum speed for 10 minutes, the supernatant was transferred to a new tube with an equal volume of isopropanol and mixed gently. Precipitation of the DNA was done at -20°C for 1h, followed by another refrigerated centrifugation at maximum speed for 10 min. The recovered DNA pellet was washed with 70% ethanol and allowed to briefly air dry. The DNA was resuspended in 1 mL of 1X TE containing RNase A to a final concentration of 100 µg/mL and incubated at 37°C for 45 min. Sodium acetate solution was added to a final concentration of 0.5 M to the RNase A-treated sample, followed by an additional chloroform extraction, isopropanol precipitation, 70% ethanol washing and brief air-drying. Finally, the DNA pellet was resuspended in ultra-pure water. The DNA quality was estimated by calculating the ratios for A260/A280 and A260/A230 using a NanoDrop spectrophotometer, and quantification was done using the Qubit fluorometer. The size and integrity of the DNA were confirmed by clamped homogeneous electrical field (CHEF) electrophoresis. For preparing genomic DNA of *Malassezia cuniculi*, we adopted two minor changes: a) the cell pellet was washed with 0.1% SDS prior to lyophilization and b) the DNA was precipitated using 100% ethanol instead of isopropanol.

**Illumina and Oxford Nanopore sequencing.** Whole-genome sequencing was performed with Illumina and Oxford Nanopore Technologies (ONT). Illumina paired-end (PE) libraries for *Malassezia* sp. CBS17886 and *Malassezia arunalokei* CBS13387 were generated and sequenced at the Duke University Center for Genomic and Computation Biology, producing 2x151-base reads. Illumina PE libraries of *Malassezia psittaci* CBS14136 and *Malassezia brasiliensis* CBS14135 were obtained and sequenced at the Genome Institute of Singapore, A*STAR, yielding 2x100-base reads. Illumina PE libraries of *Malassezia equina* CBS12830 were generated and sequenced at the Utrecht Sequencing Facility, obtaining 2x151-base reads. For nanopore sequencing, barcoded libraries were constructed as per the manufacturer's instructions using the 1D ligation sequencing kits SQK-LSK109 or SQK-LSK110 and the Native Barcoding Expansion kits EXP-NBD103, EXP-NBD104, or EXP-NBD114. DNA samples were pooled together over different MinION or PromethION R9.4.1 flow cells and sequenced in a MinION or PromethION device for 48 h. Basecalling was performed with Guppy (`-c dna_r9.4.1_450bps_hac.cfg --min_qscore=7 --qscore_filtering`) and the resulting "PASS" reads were de-multiplexed with `guppy_barcoder` (`--barcode_kits EXP-NBD104 --records_per_fastq 0 --trim_barcodes`). For *M. cuniculi* and *Malassezia caprae*, basecalled FASTQ reads from multiple runs were combined prior to assembly. See Dataset S1 for specifics on each genome.

**Genome assembly.** Genomes were assembled with ONT long-read data and polished with Illumina short reads either generated in this study or retrieved from the NCBI Sequence Read Archive (SRA) (Dataset S1). Genome assemblies were produced with Canu v2.1.1 (4) using default parameters, using genome size estimates obtained from a previous study (5). The consensus accuracy of each draft assembly was improved by first correcting errors with Medaka v1.5.0 (https://github.com/nanoporetech/medaka) or with Nanopolish v0.11.2 (https://nanopolish.readthedocs.io/en/latest/) (Dataset S1), and then with up to five rounds of polishing with Pilon v1.22 (6) (`--fix all`) with Illumina reads mapped to the first pass-polished assembly using BWA-MEM v0.7.17-r1188 (7). Contigs containing exclusively rDNA sequences detected by Barrnap (https://github.com/tseemann/barrnap) (`--kingdom euk`) or that could be assigned to mitochondrial DNA, were removed from the final assemblies. To validate telomeric regions, and to potentially identify large collapsed regions or mis-assemblies in each of the draft assemblies, Canu-corrected reads and Illumina reads were aligned with minimap2 v2.9-r720 (8) and BWA-MEM v0.7.17-r1188, respectively, and read coverage profiles were manually inspected with the Integrative Genomics Viewer (IGV) (9). No aberrant regions of coverage beyond the rDNA array region were detected. For *Malassezia dermatis* JCM11348 specifically, the available genome in GenBank (GCA_001600775.1) is assembled into 12 scaffolds. This species is closely related to *M. caprae*, with 8 chromosomes, and all species of clade B2 have conserved chromosome structures (Fig. S7). Therefore, *M. dermatis* scaffolds corresponding to rDNA or mitochondrial DNA were removed and the remaining scaffolds were re-ordered with D-GENIES (10) to match the *M. caprae* assembly. Consequently, this genome is the only one in our dataset not assembled telomere-to-telomere. Genome assemblies and primary sequencing data produced in this study were deposited at DDBJ/EMBL/GenBank (accession numbers are given in Dataset S1).

**Gene prediction and annotation.** We used funannotate v1.8.9 for gene prediction and annotation (https://github.com/nextgenusfs/funannotate) with the following steps and options. First, the contigs of each nuclear genome assembly were sorted by length (longest to shortest), labeled as chr1, chr2, etc. using `funannotate sort`, and then soft-masked for repeats with RepeatMasker v4.1.2-p1 (`-species fungi -xsmall, -nolow`). Gene prediction was performed with `funannotate predict` that calls AUGUSTUS (11), GeneMark-ES/ET (12), and EVidenceModeler (13) to generate consensus gene models. Given that no RNA-seq data was available for most species, we passed well-annotated proteins from *Malassezia sympodialis* and the UniProtKb/SwissProt curated protein database as evidence for gene prediction (`--protein_evidence`) and defined the intron length between 10 and 500 nucleotides (`--min_intronlen 10 --max_intronlen 500`) (14). For *Malassezia furfur* CBS14141, we used RNA-seq data generated in this study (described below) to train *ab initio* gene predictors using `funannotate train` (with `--jaccard_clip`, as we expected a high gene density) before running `funannotate predict`. Only the mating type gene models were inspected and manually corrected where needed (e.g., the short mating pheromone precursor gene were manually predicted and added for all species). Finally, `funannotate annotate` was run to add and integrate functional annotation to the final gene models including: PFAM and InterPro domains, Gene Ontology (GO) terms, fungal transcription factors, Cluster of Orthologous Genes (COGs), secondary metabolites, Carbohydrate-Active Enzymes (CAZYmes), secreted proteins, proteases (MEROPS), and Benchmarking Universal Single-Copy Orthologs (BUSCO) groups. InterPro data, COGs and secondary metabolite annotations were obtained by first running, respectively, InterProScan v5.55-88.0, eggNOG-mapper v.2.1.7 (eggNOG DB version: 5.0.2) (15) and AntiSMASH v6.1.0 (16), and then passing the results to *funannotate annotate (options: --iprscan, --antismash --eggnog)*.

**Ortholog identification and alignment.** To construct the phylogenomic data matrix, single-copy orthologs (SC-OGs) were identified across *Malassezia* and the outgroup *Ustilago maydis* 521 (GCA_000328475.2) using OrthoFinder v2.5.4 (17) with options: *-M msa -S diamond -I 1.5 -T iqtree -t 30 -a 4*. The amino acid sequences of 1,566 SC-OGs identified as shared among all species were individually aligned with MAFFT v7.310 (18) with arguments *--localpair --maxiterate 1000* and trimmed with TrimAl v1.4.rev22 (19) using *-gappyout -keepheader*.

**Phylogenetic analyses and topological support evaluation.** The evolutionary relationships within *Malassezia* were inferred using two different approaches: (i) concatenation with gene-based partitioning in IQ-TREE v2.1.3 (20), and (ii) gene-based coalescence in ASTRAL-MP v5.15.2 (21). For the first method, a maximum likelihood (ML) phylogenetic tree we inferred based on individual protein alignments passed to IQ-TREE with the argument *"-p"*. This concatenates all alignment files into a supermatrix (21 taxa, with 1,566 partitions and 950,691 sites) for partition analysis and applies an edge-linked proportional partition model to accommodate different evolutionary rates between partitions (22). The best-fitting amino acid substitution model for each partition was determined with ModelFinder (23) and the Bayesian information criterion (BIC). The best-scoring ML tree was inferred using the parameters *--seed 54321 -m MFP -msub nuclear -B 1000 -alrt 1000 -T 20*, which also assesses support for each internal branch using 1,000 replicates of the Shimodaira–Hasegawa approximate likelihood ratio test (SH-aLRT) and ultrafast bootstrap (UFboot) (24). For the second method, we first inferred ML gene trees for each of the 1,566 SC-OGs with IQ-TREE using the argument *"-S"* that performs model selection and tree inference for each alignment individually. The coalescent-based species phylogeny was inferred with ASTRAL using default settings and a file containing 1,566 individual ML gene trees as input. The option *"-t 2"* was used to retrieve the full set of different measurements, including quartet support for the main (q1) and alternative topologies (q2 and q3), and the local posterior probability (LPP) support values. Genealogical concordance and topological support metrics were analyzed for each bipartition of the best-scoring ML phylogeny by measuring the gene concordance factor (gCF) and the site concordance factor (sCF) as implemented in IQ-TREE (25). The gCF describes the fraction of single-gene trees that can recover a particular branch, and sCF is defined as the percentage of sites of the alignment that provide decisive support for a given branch in the tree. To evaluate both measures within a single run, the file containing the best-scoring ML tree derived from gene-based concatenation (*concat.treefile*) and the file containing 1,566 single-gene trees (*Loci.treefile*) were input to IQ-TREE with the options: *-t <concat.treefile> --gcf <Loci.treefile> -p <Alignment_dir> --scf 100*. All the trees were visualized in iTOL v5.6.3 (26).

**Mating-type genes and gene genealogies.** *MAT* regions in *Malassezia* genomes were identified by BLAST searches using the well-annotated *MAT*-derived proteins from *M. sympodialis* (27), and manually reannotated if necessary. The short pheromone precursor genes were not among the automatically predicted genes and were instead identified manually by translating the regions flanking the pheromone receptor genes and retrieving the product that ended with a C-terminal CAAX motif characteristic of fungal mating pheromones (28) (Fig. 2C). Pheromone receptor protein alignments were generated and trimmed as above. Predicted lipases (i.e., protein sequences containing LIP family PF03583 or Class 3 family PF01764 domains; Figs. S16 and S17) and secretory aspartyl proteases (containing the PF00026 domain; Figure S18) were extracted from the corresponding orthogroups as determined by Orthofinder, aligned, and trimmed as above. The phylogeny of *M. furfur* and *M. brasiliensis HD* alleles (Fig. S13) was obtained by first retrieving, aligning, and trimming the sequences encompassing the complete region of the *bW* and *bE* genes, including their common intergenic region. Maximum likelihood (ML) phylogenies were inferred with IQ-TREE2 and visualized with iTOL v5.6.3. The specific model parameters for phylogenetic reconstruction are given in the respective figure legends.

**GC-content analysis and centromere identification.** The genome-wide GC-content was determined with SeqKit v2.0.0 (29) using non-overlap windows of 0.25 kb and 1 kb (`seqkit sliding -s 250 -W 250 | seqkit fx2tab -n -g -i`, adjusting parameters `-s` and `-W` accordingly). The 1 kb-window data were plotted as distributions across species using ggplot2 v3.4.1 with ggridges v0.5.4 in R v4.2.1 (ridgeline plots, Fig. 1C), whereas the 0.25 kb-window data were plotted along the chromosomes of each species, together with the locations of candidate centromeres and mating-type loci (Fig. S3) using the Bioconductor karyoploteR v1.22 R package. The local GC-content bias was further evaluated with OcculterCut (30) with default settings, which segments the genome sequences into regions of differing GC-content using the Jensen–Shannon divergence and identifies AT-rich and GC-equilibrated regions (Fig. S3). *Malassezia* centromeres map to chromosomal GC-poor troughs, but their exact location is difficult to pinpoint given their reduced size and the fact they may overlap with active genes located proximal to these troughs (31). Therefore, the location of the centromeres in the new assemblies was determined by inspecting the GC-content in sliding windows of 250 bp (step size = 2) and retrieving the coordinates (given in Dataset S1) corresponding to the regions on each side of the GC-minimum where GC% values are lower than 40%.

**Identification of telomeres and survey for telomere-adjacent repeats.** Telomeric repeats were identified by manually inspecting contig ends in all genomes. Motifs that were tandemly repeated at contig termini were assigned as candidate telomeric repeat sequences. The accuracy of the assembly and consensus sequences at these regions was validated by visual inspection in IGV after aligning the respective Canu-corrected and Illumina reads to the assembly with minimap2 v2.9-r720 and BWA-MEM v0.7.17-r1188, respectively. Four different motifs were identified across species: TAACCC (in *M. cuniculi*, *Malassezia slooffiae*, *Malassezia* sp. CBS17886 and *Malassezia vespertilionis*); TTAACAC (in *Malassezia nana*, *M. equina*, *M. sympodialis* and *M. caprae*); TAACAC (in *Malassezia pachydermatis*, *Malassezia globosa*, *Malassezia restricta* and *M. arunalokei*); and TAATCC (in *M. japonica*, *M. obtusa*, *M. psittaci*, *Malassezia yamatoensis*, *M. brasiliensis*, *M. furfur* P1 and *M. furfur* P2) (summarized in Table S1). Additionally, contigs were inspected for inverted repeat regions using UGENE v40 (32) with window size = 100 bp and minimum identity per window = 95%. Inverted repeats found at both ends of the same contig extending inwards from the telomere repeat tract were further analyzed. First, telomere repeats were stripped off to generate a set of telomere-adjacent sequences. Then, the resulting sequences were translated, and the amino acid sequence corresponding to the longest predicted open read frame (ORF) was analyzed for protein domains using CD-Search (NCBI) and InterProScan (EMBL-EBI) and used as queries in BLASTP searches. A reverse transcriptase domain (PF00078) was identified in all sequences, and BLASTP produced top matches to hypothetical proteins of other basidiomycetes, including the *Cryptococcus neoformans* Cnl1 non-long-terminal repeat (non-LTR) element (sequences are provided in Dataset S1). Telomeric repeats, inverted repeats, and candidate non-LTR elements were plotted using Bioconductor karyoploteR v1.22 with GenomicFeatures v1.48.4 R packages and edited in Adobe Illustrator to improve figure rendering (Fig. S2).

**Synteny analyses.** Synteny relationships between all possible pairs of *Malassezia* genomes were determined using SynChro (33) included within the CHROnicle package (http://www.lcqb.upmc.fr/CHROnicle/) setting the block stringency (delta) parameter to 3. SynChro uses a file containing a list of ordered genes and their amino-acid sequences as input, which was obtained by parsing the GFF3 and protein output files of the funannotate pipeline using standard text-parsing Unix tools. The *M. sympodialis* genome was used as a reference in our comparisons. Linear synteny plots comparing chromosomes and specific genomic loci (e.g., mating-type loci regions) were generated with EasyFig (34) by parsing and splitting the GenBank (.gbk) output files of the funannotate pipeline using standard text-parsing Unix tools. EasyFig plots were produced used the following general parameters: `-blastn -svg -f gene frame -legend both -leg_name locus_tag -blast_col 180 215 242 180 215 242 -blast_col_inv 249 176 154 249 176 154 -bo F -width 4000 -ann_height 100 -blast_height 200 -f1 T -f2 10000 -G GCContent -wind_size 100 -step 100 -g_height 200 -line T`. Color codes and labels in the SynChro and EasyFig plots were edited in Adobe Illustrator for figure rendering.

**Fusion and mating assays.** Initially, fusion experiments were conducted for *M. furfur* and *M. sympodialis.* Equal amounts of *M. furfur ade2*Δ mutants generated in strain CBS14141 (*MAT a2b1*) (2) and *M. furfur* CBS9369 (*MAT a1b2*) (35), and *M. sympodialis* ATCC42132 (*MAT a1b1*) resistant to nourseothricin ($NAT^R$) and ATCC44340 (*MAT a2b2*) resistant to neomycin G418 ($NEO^R$) (2), were mixed or spotted alone on different culture media (YPD, YT, Filament agar, YNB, MS, PDA, V8 pH5, V8 pH7) and incubated for 2 days at 30°C. A modified version of these media (hereafter designated as modified or mYPD, mYT, etc.)

that contained 1 mL/L of Tween 20, 4 mL/L of Tween 60, and 4 g/L of Ox Bile to fulfill *Malassezia* lipid requirements and improve growth was also tested. Both the mix and single spots of each strain were resuspended in sterile water and plated on modified Minimal Medium (mMM) + nourseothricin (100 µg/mL) for *M. furfur*, and on mDixon + nourseothricin (100 µg/mL) and neomycin G418 (100 µg/mL) for *M. sympodialis.* Afterwards, a larger collection of *M. furfur* isolates was obtained (Table S2). All these *M. furfur* strains were subjected to mating assays by co-culturing equal proportion of single strains onto the different media indicated above and under a variety of growth conditions (dark, light, room temperature, 30°C). Last, the addition of several compounds [squalene (1ml/L), potassium nitrate (1 g/L), sodium chloride (1.3 g/L), ferrous sulphate (0.01 g/L), magnesium sulphate (0.15 g/L) and glycine (3.75 g/L)] previously reported to induce filamentation in *Malassezia* was also tested (36); media containing these compounds are reported in the text as "+ supplements" (e.g., YPD + supplements). Plates were monitored by microscopy for a period of 20 to 40 days.

***Agrobacterium tumefaciens*-mediated transformation.** A region of 2,323 bp including the mating pheromone and pheromone receptor genes of the *MAT a1* (*P/R*) allele of *M. furfur* strain CBS9369 (*MAT a1b2*) (35) was amplified by PCR using primers JOHE43874 and JOHE43875. The *NEO* marker was obtained from plasmid pAIM5 (2) with primers ALID2078 and ALID2081. A region of 3,850 bp including the *bE* and *bW* genes of the *MAT b4* (*HD*) allele was amplified from the diploid *M. furfur* strain CBS7710 (5) with primers JOHE43876 and JOHE43877. Primers JOHE43874 and JOHE43877 have chimeric regions complementary to the binary vector pGI3 to allow recombination in *S. cerevisiae*, while primers JOHE43875 and JOHE43876 have chimeric regions complementary to the *NEO* marker. The three PCR products, including the *MAT a1* allele, the *NEO* marker, and the *b4* allele, were assembled within the T-DNA region of plasmid pGI3 linearized with BamHI and KpnI through *in vivo* recombination in *S. cerevisiae* as previously described (2). *S. cerevisiae* transformants containing the newly recombined *a1-NEO-b4* plasmid (named pGI26), were identified by colony PCR using primer pairs JOHE43279-JOHE43280 and JOHE43281-JOHE43282 to assess correct recombination at the 5′ and 3′ junctions, respectively. Positive transformants were subjected to DNA extraction using phenol:chloroform:isoamyl alcohol 25:24:1. The extracted DNA was used to transform *A. tumefaciens* EHA105 through electroporation, with selection on LB + Kanamycin 50 µg/mL. Subsequently, plasmid pGI26 was extracted from *A. tumefaciens* and used to generate two other plasmids: one that included only the *a1-NEO* region, and another that only contained the *NEO-b4* region. Specifically, the *a1-NEO* region was amplified from plasmid pGI26 using primer pairs JOHE43874-JOHE 44551, whereas the *NEO-b4* region was amplified using primers JOHE46235-JOHE43877. These regions were independently inserted within the T-DNA of plasmid pGI3 through *in vivo* recombination as previously described, generating plasmids pGI52 and pGI55, respectively. PCR reactions were carried out with Phusion High Fidelity DNA polymerase (New England Biolabs) and consisted of 34 cycles of denaturation at 98°C for 30 sec, annealing at 60°C for 30 sec, extension at 72°C of 1 min, with an initial denaturation at 98°C for 2 min and a final extension at 72°C for 5 min. *Agrobacterium*-mediated transformation was performed as previously described (37). Briefly, *M. furfur* strain CBS14141 was incubated for 2 days at 30˚C in mDixon liquid medium, and *A. tumefaciens* strains containing plasmids pGI26 (*a1-NEO-b4*), pGI52 (*a1-NEO*) and pGI55 (*NEO-b4*) were incubated overnight in LB + kanamycin at 30°C. Both *M. furfur* and *A. tumefaciens* stains were diluted to an $OD_{600}$ of 1 and mixed at 2:1, 5:1 and 10:1 *Malassezia*:*Agrobacterium* ratios and pelleted at 5000 rpm for 10 min. The resulting cell pellet was spotted onto nylon membranes on modified Induction medium (mIM) plates and incubated at 24°C for 3 days. Cells were scraped off the membranes into sterile $H_2O$ and pelleted at 2700 rpm for 5 min, and the resulting pellets were resuspended in sterile $H_2O$ and spread onto mDixon with neomycin G418 (100 µg/mL) and cefotaxime (300 µg/mL). Resulting colonies were screened by PCR to confirm the ectopic presence of the inserted T-DNA molecules using primers specific for the *MAT* regions, such as JOHE44272-JOHE44273 for the *MAT a1*, JOHE44274-JOHE44275 for *MAT a2*, JOHE44491-JOHE44492 for the *MAT b1*, and JOHE44494-JOHE44496 for the *MAT b2* (35); JOHE43965-JOHE43966 were used for the *NEO* marker. All primers used in this study are listed in the Table S3.

**Light microscopy.** *M. furfur* WT CBS14141 and selected transformants were inoculated in solid media in the same conditions as reported in the section "Fusion and mating assays". Cells were imaged with a Nikon Eclipse E400 microscope equipped with a Nikon DXM1200F camera directly on plates or after transferring them onto glass slides. For analysis of hyphal growth in liquid culture, *M. furfur* WT CBS14141 and selected transformants were inoculated in liquid YPD + supplements at 32°C for several days. On a daily basis, an aliquot of each sample was withdrawn and subjected to microscopy analysis using a Zeiss Imager widefield fluorescence microscope.

**Scanning electron microscopy (SEM).** *M. furfur* WT CBS14141 and selected transformants were grown overnight in mYPD at 32°C. Several aliquots of 25 µL of these cellular suspensions were spotted onto YPD agar + supplements and kept at 30°C for 8 days. Samples were prepared for SEM as previously described (38). Briefly, several agar blocks containing hyphae were excised and fixed in 0.1 M sodium cacodylate buffer, pH = 6.8, containing 3% glutaraldehyde at 4°C for several weeks. Before imaging, the agar block was rinsed with cold 0.1 M sodium cacodylate buffer, pH = 6.8 three times and post-fixed in 2% osmium tetroxide in cold 0.1 M cacodylate buffer, pH = 6.8 for 2.5 hours at 4°C. Then the block was critical-point dried with liquid $CO_2$ and sputter coated with 50 Å of gold/palladium with a Hummer 6.2 sputter coater (Anatech). The samples were viewed at 15KV with a JSM 5900LV scanning electron microscope (JEOL) and captured with a Digital Scan Generator (JEOL) image acquisition system. SEM was performed at the North Carolina State University Center for Electron Microscopy, Raleigh, North Carolina, USA.

**RNA-sequencing.** *M. furfur* WT CBS14141 and derived transformants were grown overnight in 10 mL of mYPD. Five mL of these cultures were inoculated onto 100 mL of YPD + supplements. Cultures were grown at 30°C for 2 days and evaluated under the microscope for the presence of hyphae. Cells were centrifuged at 5,000 rpm for 3 minutes, the pellet washed with 5 mL of $dH_2O$, lyophilized, and frozen at -80°C. RNA extraction was performed using TRIzol (ThermoFisher) as previously described (39). Final RNA pellets were treated with TURBO DNase (ThermoFisher) according to manufacturer's instructions, and RNA was resuspended in 50 µL of nuclease free water (Promega). Illumina Single End (SE) libraries were prepared using the TruSeq Stranded Total RNA-seq Kit and sequenced on an Illumina HiSeq 2500 platform to generate SE 50-base reads. Library preparation and RNA sequencing were performed at the Duke University Center for Genomic and Computation Biology. Three biological replicates were performed and sequenced for each sample.

**Differential expression analysis.** Raw SE reads were filtered and trimmed with Trim Galore v0.6.7 using default settings (phred score = Q20) (https://github.com/FelixKrueger/TrimGalore). Reads that became shorter than 45 nt, because of quality or adapter trimming, were discarded (option: `--length 45`). The remaining reads were mapped to a reference genome comprising the *M. furfur* CBS14141 nuclear genome assembly plus an added contig corresponding to the sequence of the construct containing both the *P/R* and

*HD* loci used for transformation (i.e., the *A*on *B*on construct). This approach allows verifying if the introduced alleles, which differ from those of the parental strain, are expressed in the *A*on and *A*on *B*on strains. Read mapping was performed with STAR v.2.7.4a (40) using default settings, except that the maximal intron length was reduced to 200 bp. The resulting alignment BAM files were visually inspected with IGV v2.14. Read summarization and count matrices were obtained from the alignment files with featureCounts v2.0.1 (41) and differential expression between *M. furfur* transformants (*A*on or *A*on *B*on) and the parental strain (WT control) was analyzed using the Bioconductor DESeq2 v1.36.0 R package, which report *p*-values that indicate statistical significance, i.e., likelihoods that a gene is not differentially expressed (42). For each comparison (*A*on vs. WT, and *A*on *B*on vs. WT), genes with a false discovery rate (FDR) < 0.05 (*p*-value, adjusted for multiple testing with the Benjamini-Hochberg procedure) and $\log_2$ fold change (FC) > ± 1.0, were considered differentially expressed. To assess between sample variability hierarchical clustering of gene expression level profiles of the different samples was plotted with regularized-logarithm (rlog) transformation of DESeq2-normalized expression data. Volcano plots showing gene expression between pairs of samples were generated using the Bioconductor EnhancedVolcano v1.14.0 (https://github.com/kevinblighe/EnhancedVolcano) R package from the log2 FC of all detected transcripts. Differential expression data is provided as Datasets S1, S2 and S3.

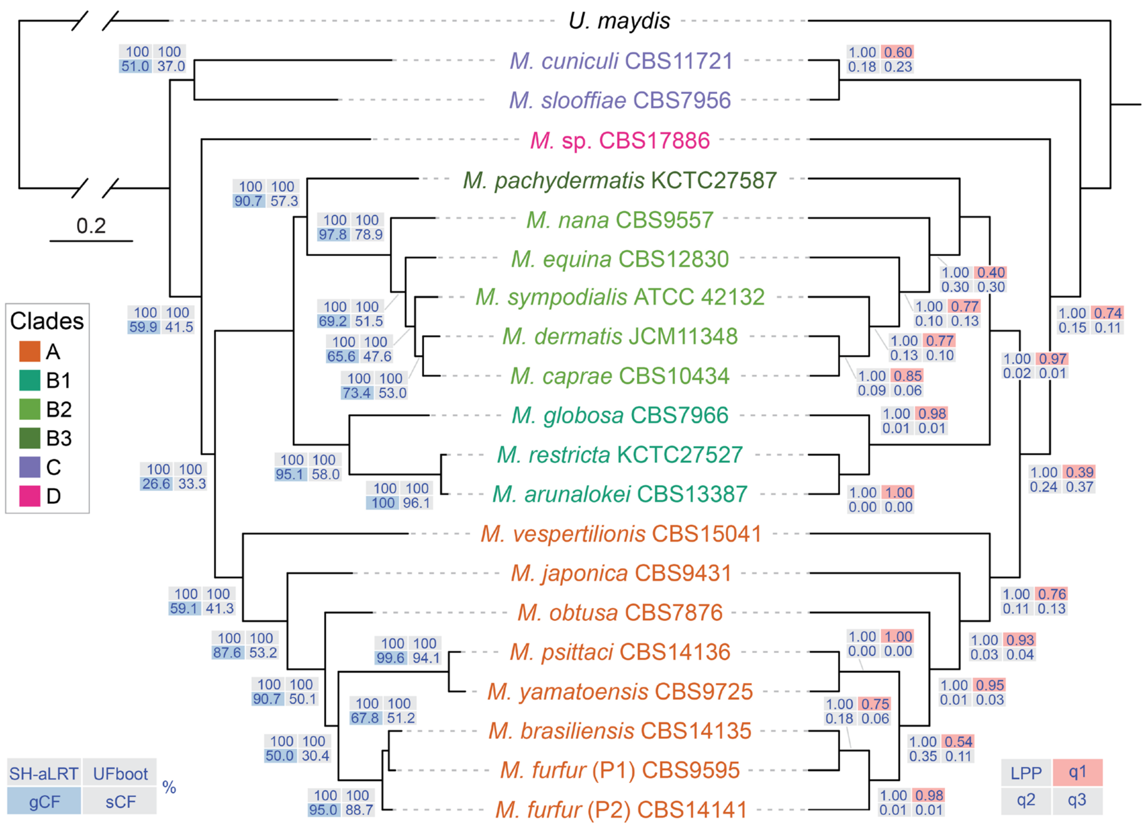


**Fig. S1.** Tree topologies inferred by concatenation- and coalescence-based approaches. The maximum likelihood (ML) tree derived from a concatenation-based approach (left) is fully congruent with the tree topology derived from the coalescence approach (right). For the coalescence analysis, all branches received 1.0 local posterior probability (LPP) and the other numerical values on the branches represent quartet support for the main topology (q1) and alternative topologies (q2 and q3). For the ML phylogeny, support values were obtained from 1000 replicates of the Shimodaira–Hasegawa approximate likelihood ratio test (SH-aLRT) and ultrafast bootstrap (UFboot), and the additional numeric values represent the proportion of genes (gCF) or sites (sCF) that are concordant with a given branch in the tree.

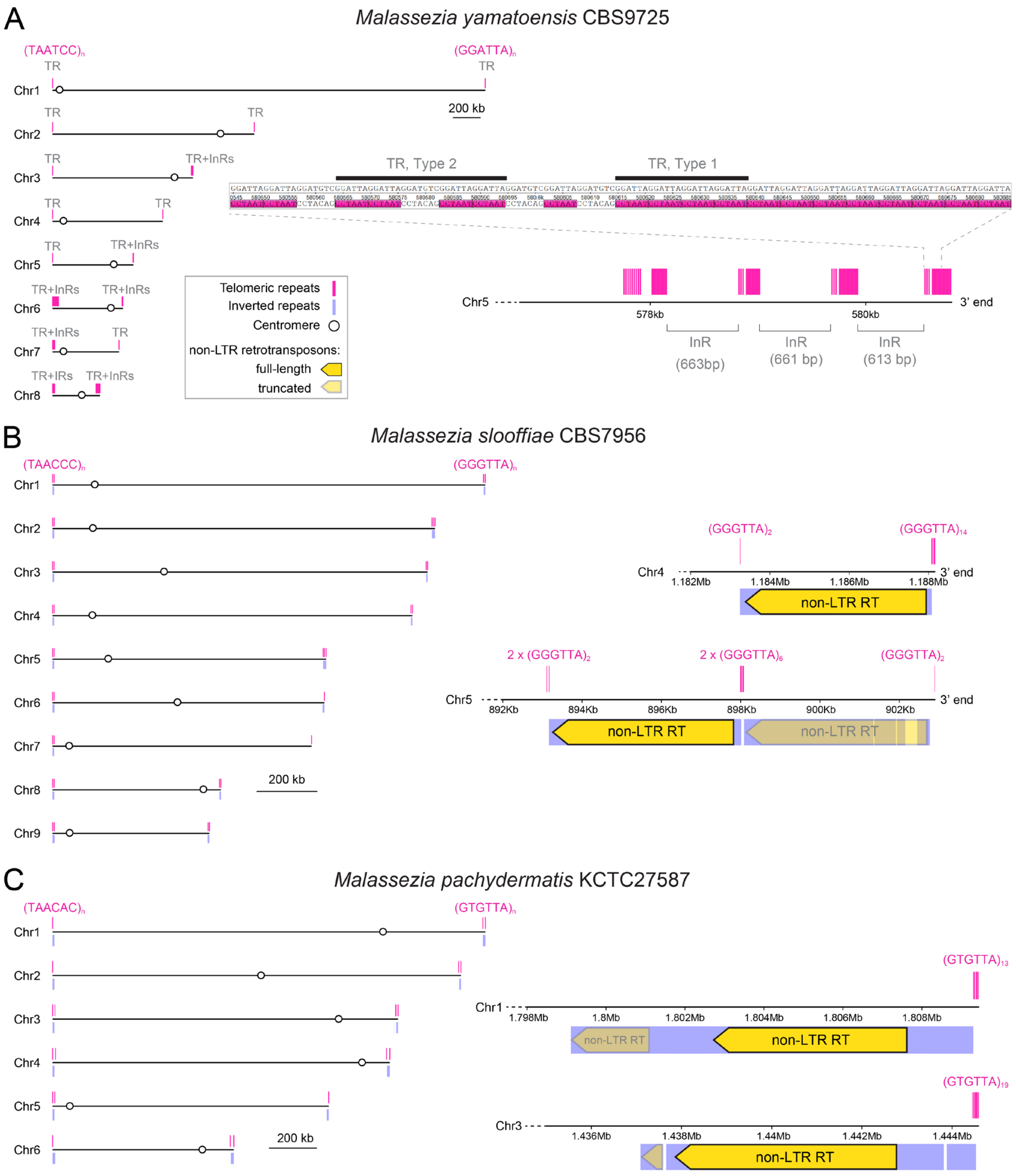


**Fig. S2.** Chromosome ends in *Malassezia*. (A) In *M. yamatoensis*, 7 out of 16 chromosome ends are composed of blocks of telomeric repeat sequences interspersed with unknown repeat sequences (TR+InRs). These blocks extend over several kb in some cases (e.g., at the left end of chr. 6 and right end of chr. 8) and encompass two major repeat patterns (TR, types 1 and 2). (B and C) In *M. slooffiae* and species of sub-clades B2 and B3 (*M. pachydermatis* is illustrated here as an example), chromosome ends contain inverted repeats predicted to encode non-long-terminal repeat (non-LTR) retrotransposons (RT) followed by the terminal telomeric repeat tract.

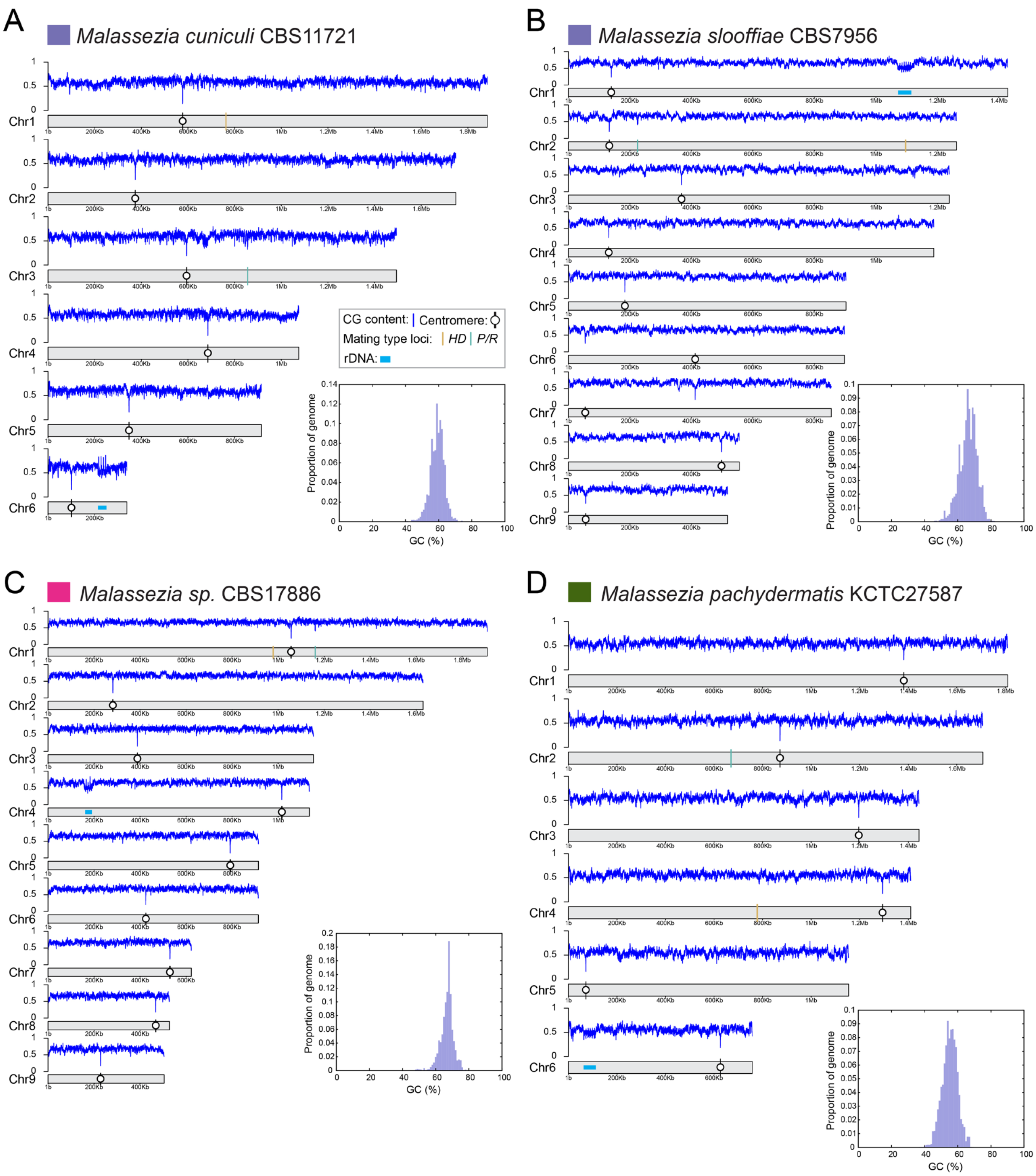


**Fig. S3.** Plots depicting chromosomes and locations of centromeres, rDNA and *MAT* loci. The GC content is plotted both along each contig and as a density plot for the whole-genome. All four species present a unimodal pattern of GC-content distribution as determined by OcculterCut (30).

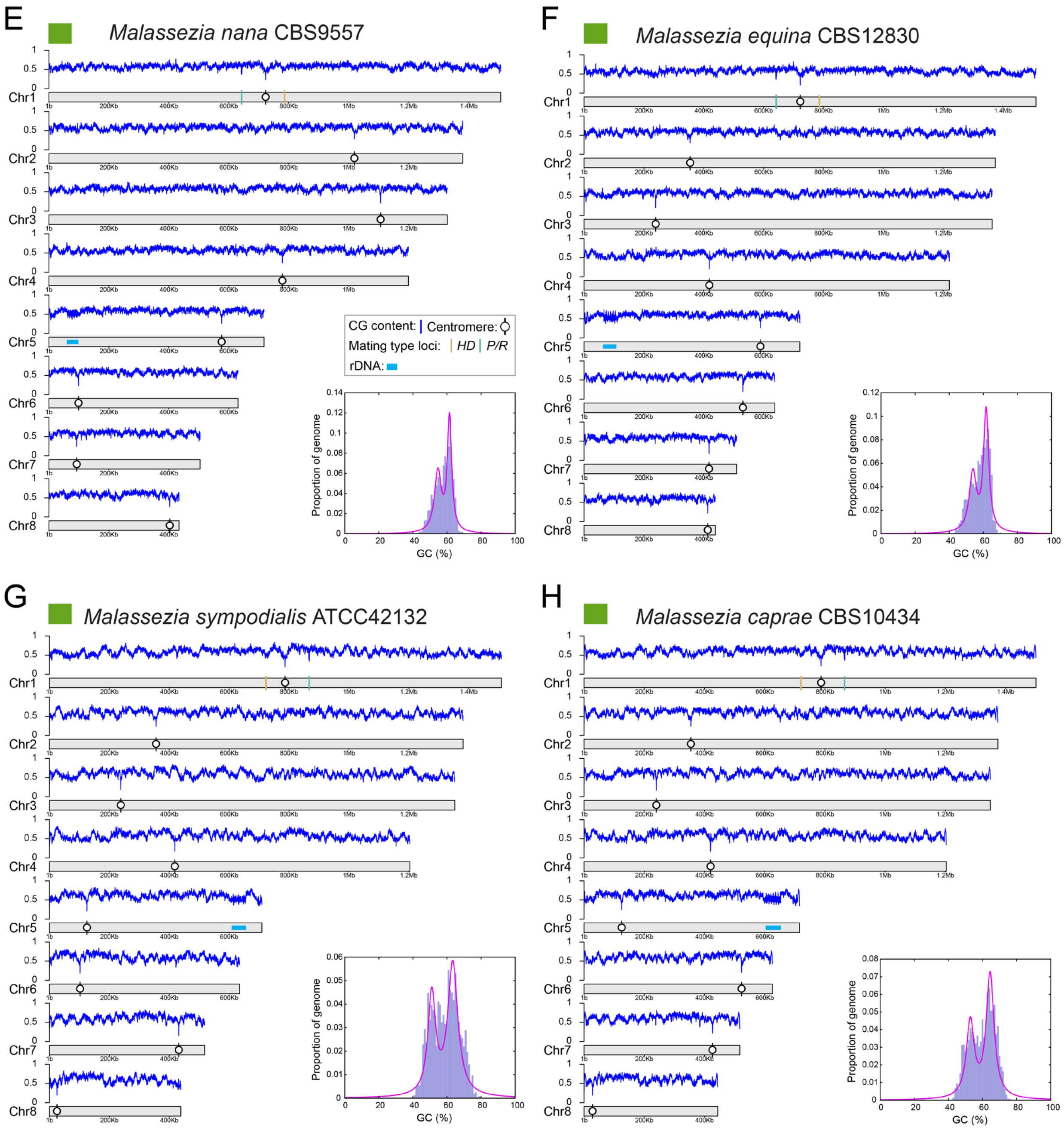


**Fig. S3.** (continued). Plots depicting chromosomes and locations of centromeres, rDNA and *MAT* loci. The GC content is plotted both along each contig and as a density plot for the whole-genome. All four species present a broader and partially bimodal pattern of GC-content distribution as determined by OcculterCut (30).

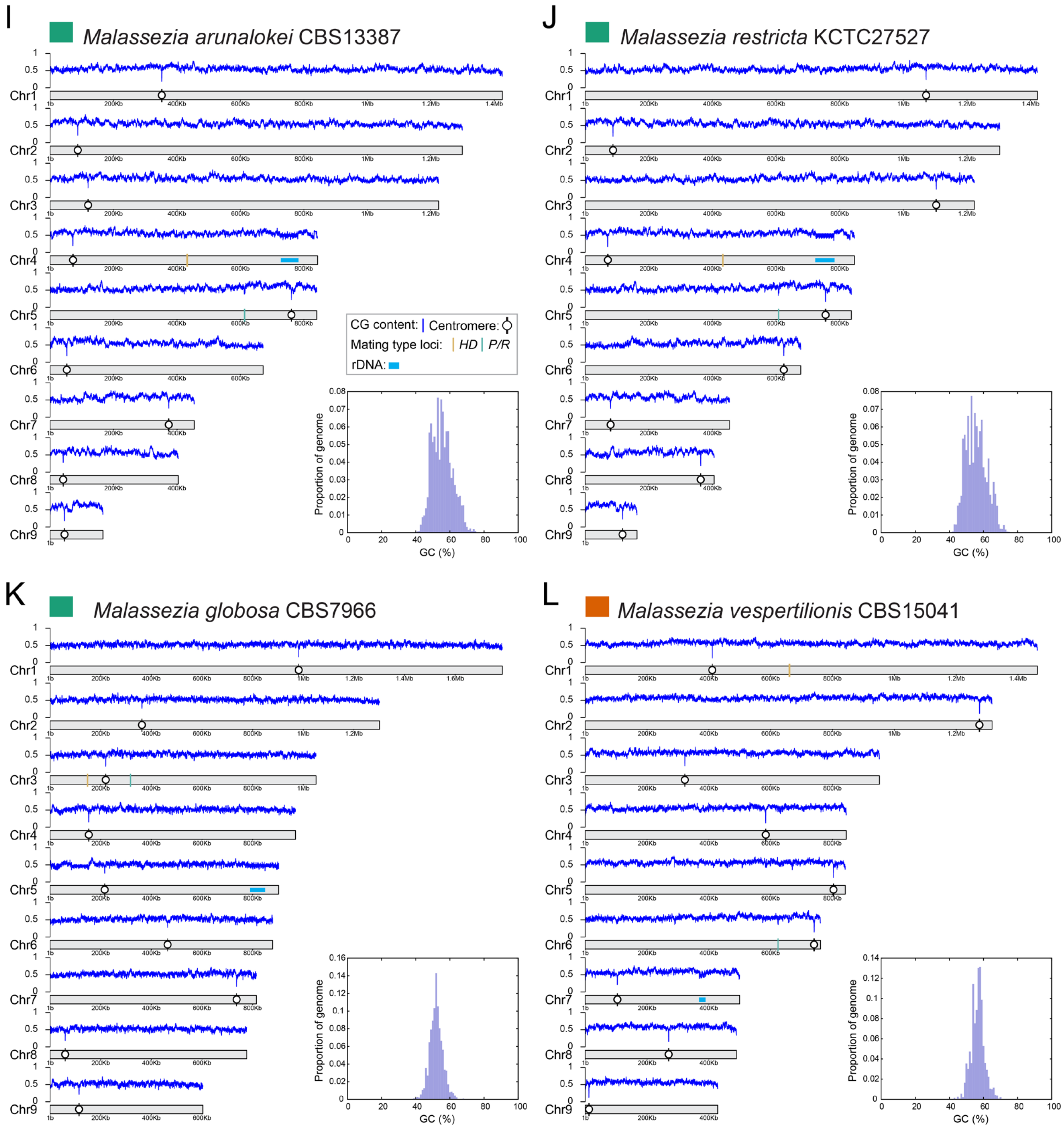


**Fig. S3.** (continued). Plots depicting chromosomes and locations of centromeres, rDNA and *MAT* loci. The GC content is plotted both along each contig and as a density plot for the whole-genome. All four species present a unimodal pattern of GC-content distribution as determined by OcculterCut (30).

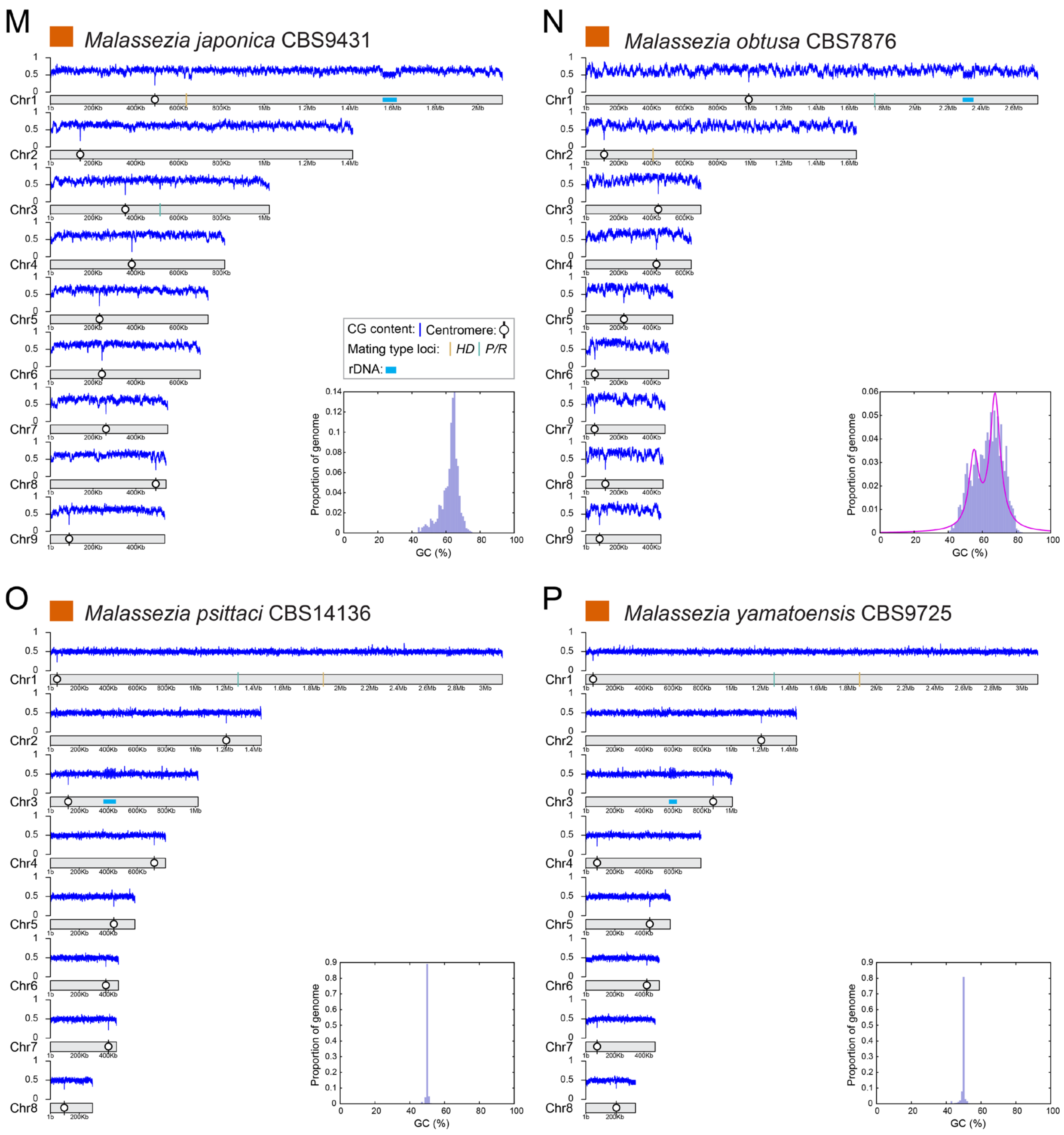


**Fig. S3.** (continued). Plots depicting chromosomes and locations of centromeres, rDNA and *MAT* loci. The GC content is plotted both along each contig and as a density plot for the whole-genome. All of these species present a unimodal pattern of GC-content distribution as determined by OcculterCut (30), except *M. obtusa* that presents a broader and partially bimodal profile. The genomes of *M. psittaci* and *M. yamatoensis* have the lowest GC-content and show an extremely narrow GC-content distribution.

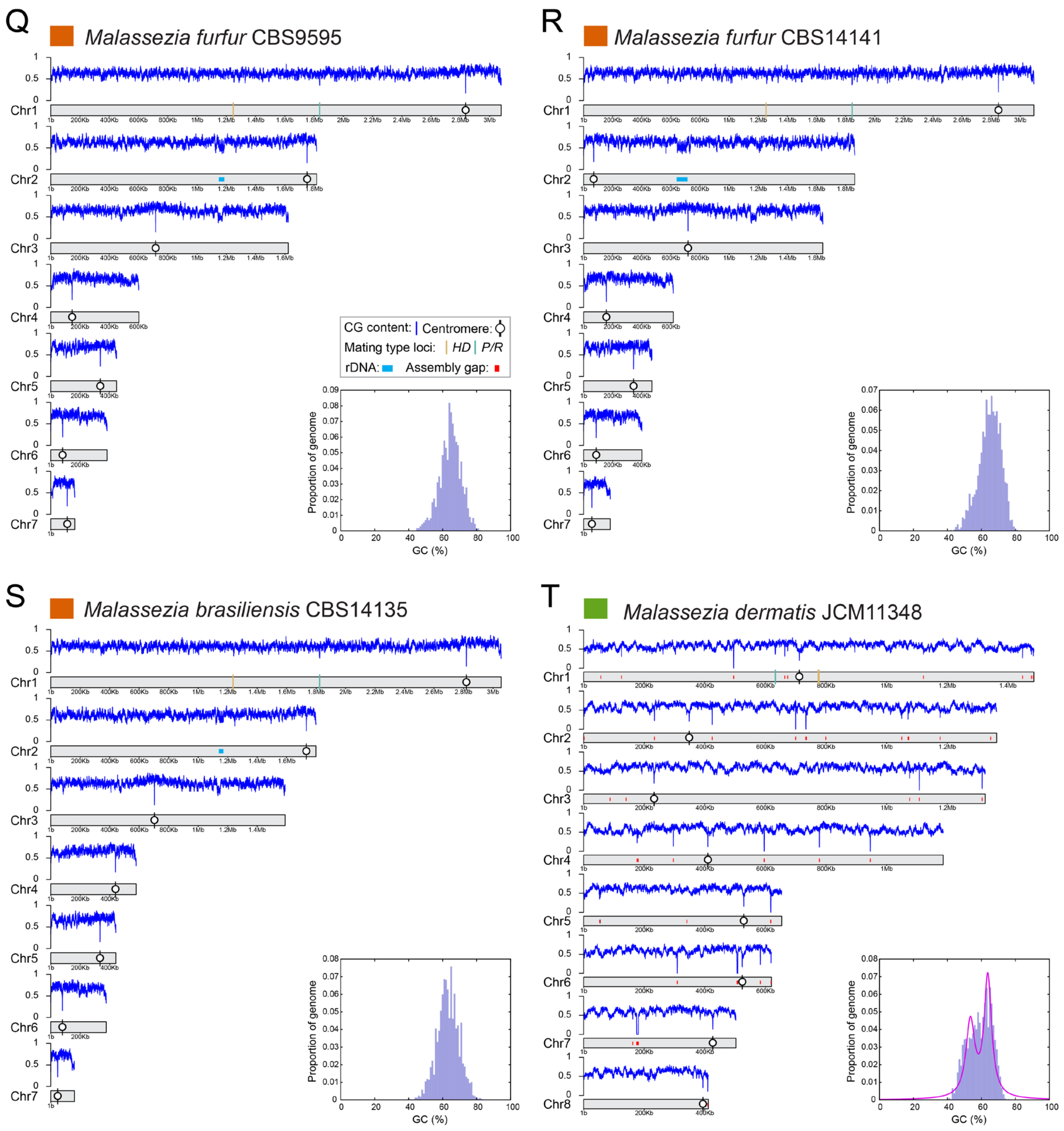


**Fig. S3.** (continued). Plots depicting chromosomes and locations of centromeres, rDNA and *MAT* loci. The GC content is plotted along each contig and as a density plot for the whole-genome. All these species, expect *M. dermatis*, present a unimodal pattern of GC-content distribution as determined by OcculterCut (30). The genome assembly of *M. dermatis* was not re-sequenced in this study and still contains a few gaps (depicted by red dashes).

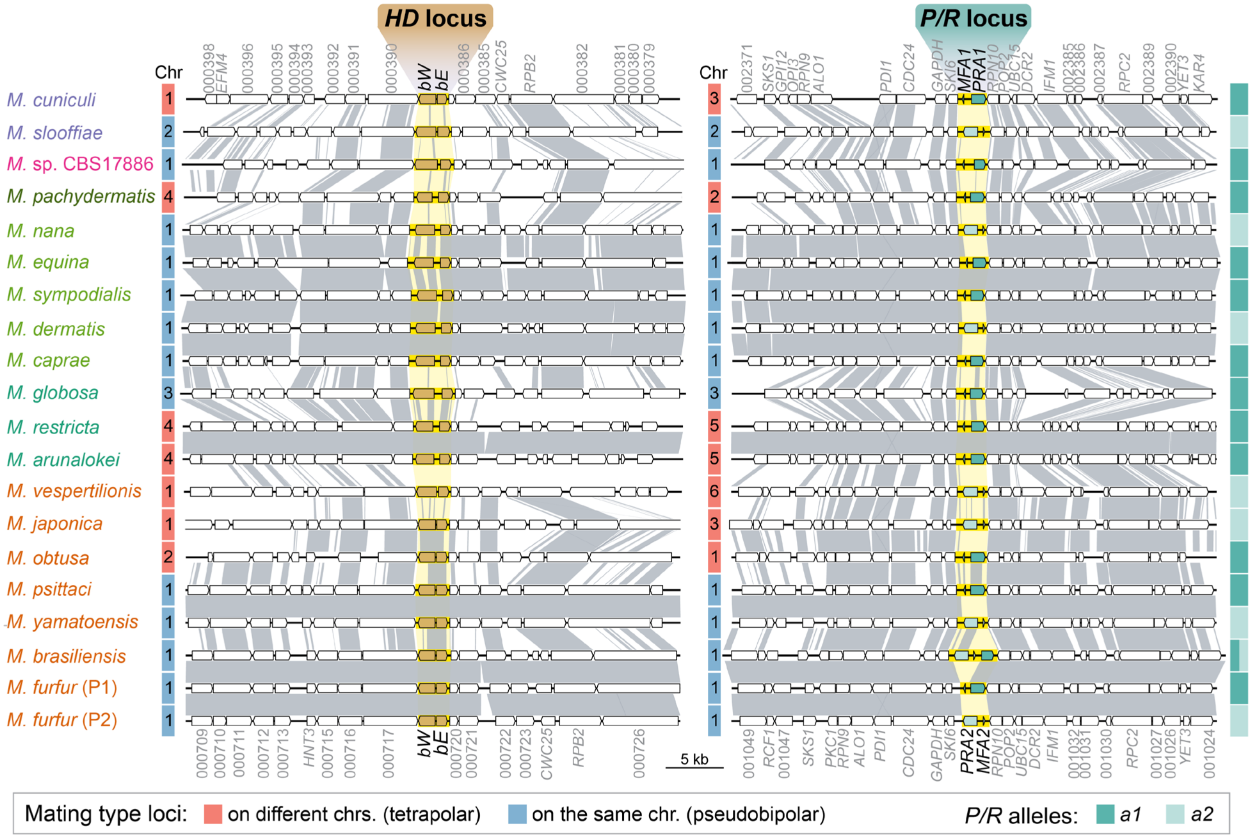


**Fig. S4. Mating-type (*MAT*) loci structure and *MAT* genes in *Malassezia*.** (A) Synteny analysis comparing the chromosomal regions encompassing the *HD* and *P/R MAT* loci regions across all species. The *P/R* and *HD* loci are highlighted in yellow. The *HD* locus comprises two divergently transcribed genes *bE* (HD1-type) and *bW* (HD2-type). The *P/R* locus comprises only one pheromone receptor gene (*PRA*) and one pheromone precursor gene (*MFA*), except in *M. brasiliensis*, which encodes two alternate pheromone receptors (Pra1 and Pra2), but only one pheromone precursor (Mfa2). In pseudobipolar species the *P/R* and *HD* loci are located on the same chromosome (Chr) at variable distances, whereas in tetrapolar species, the two *MAT* loci reside on different chromosomes.

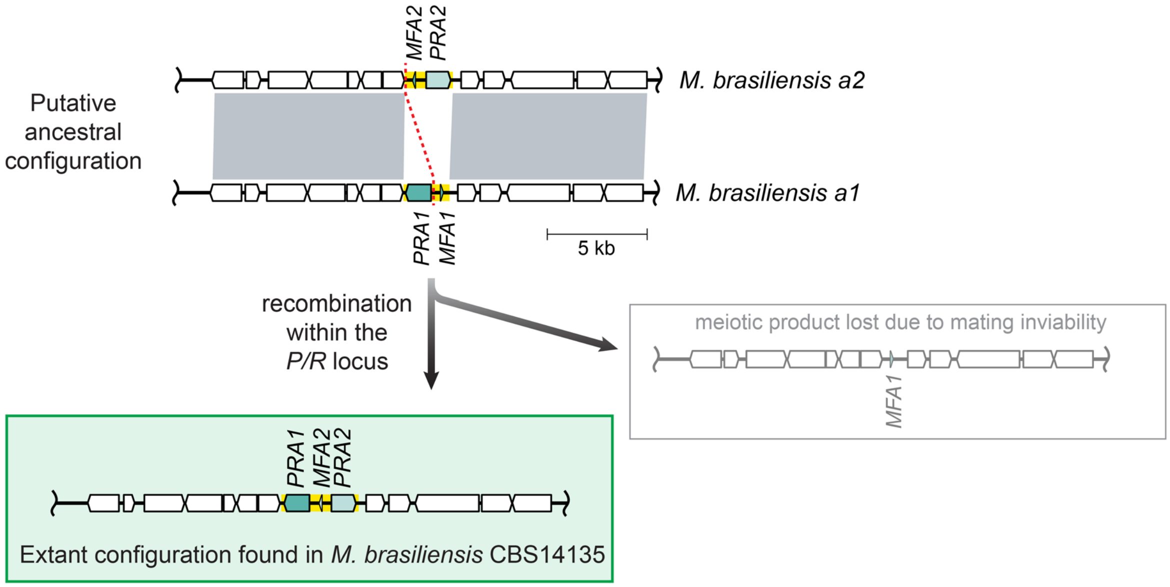


**Fig. S5.** The *P/R* locus of *M. brasiliensis* likely originated via recombination between opposite mating types. Recombination at the *P/R* locus probably involved the common intergenic region of *PRA1* and *MFA1* in the *a1* allele and the downstream region of *MFA2* in the *a2* allele. Such an event would produce a recombinant progeny containing a *P/R* locus with two alternate receptors and one pheromone gene, and another product harboring only the other pheromone gene. The latter product was likely lost due to mating inviability as it is unable to sense the pheromone produced by the opposite mating type. Conversely, the other product would be able to sense and produce mating pheromone allowing intercommunication with a compatible mating partner. No repetitive sequences were found at the candidate cross-over breakpoint.

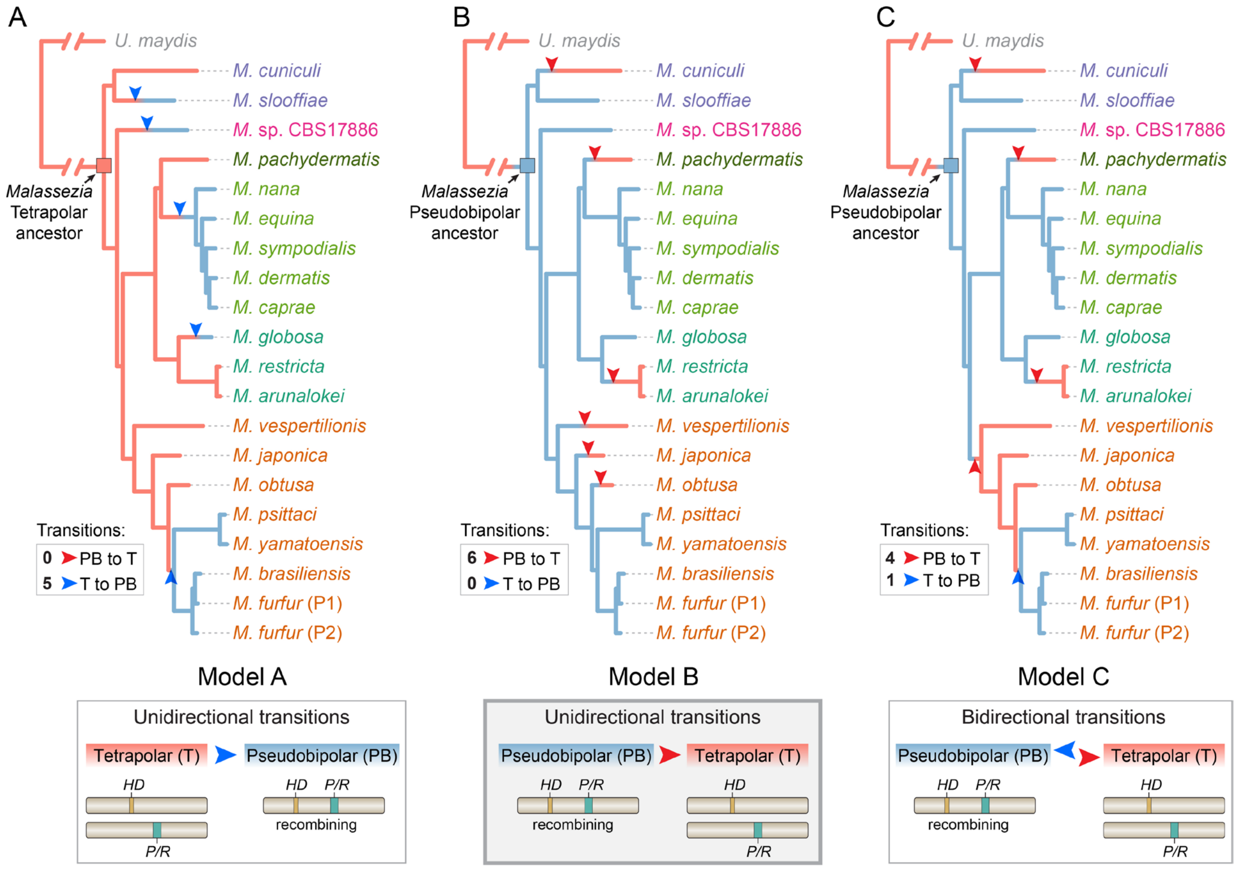


**Fig. S6.** Hypotheses on the evolution of mating-type loci chromosomal organization in *Malassezia*. Orange and blue branches depict, respectively tetrapolar and pseudobipolar configurations. (A) Model A assumes unidirectional transitions and the presence of a tetrapolar configuration in the most recent common ancestor of *Malassezia*. Under this assumption, five independent transitions to pseudobipolar (blue arrowheads) would have had to occur to explain the extant *MAT* loci organization across species. (B) If the *Malassezia* ancestor was instead pseudobipolar, six independent unidirectional transitions to tetrapolarity (red arrowheads) would have occurred during evolution (model B). Finally, assuming a pseudobipolar ancestral configuration and considering that transitions can be bidirectional, four independent pseudobipolar to tetrapolar transitions are hypothesized with a possible reversal to pseudobipolarity in one case. The analyses presented in this study supports model B (refer to Fig. 3A).

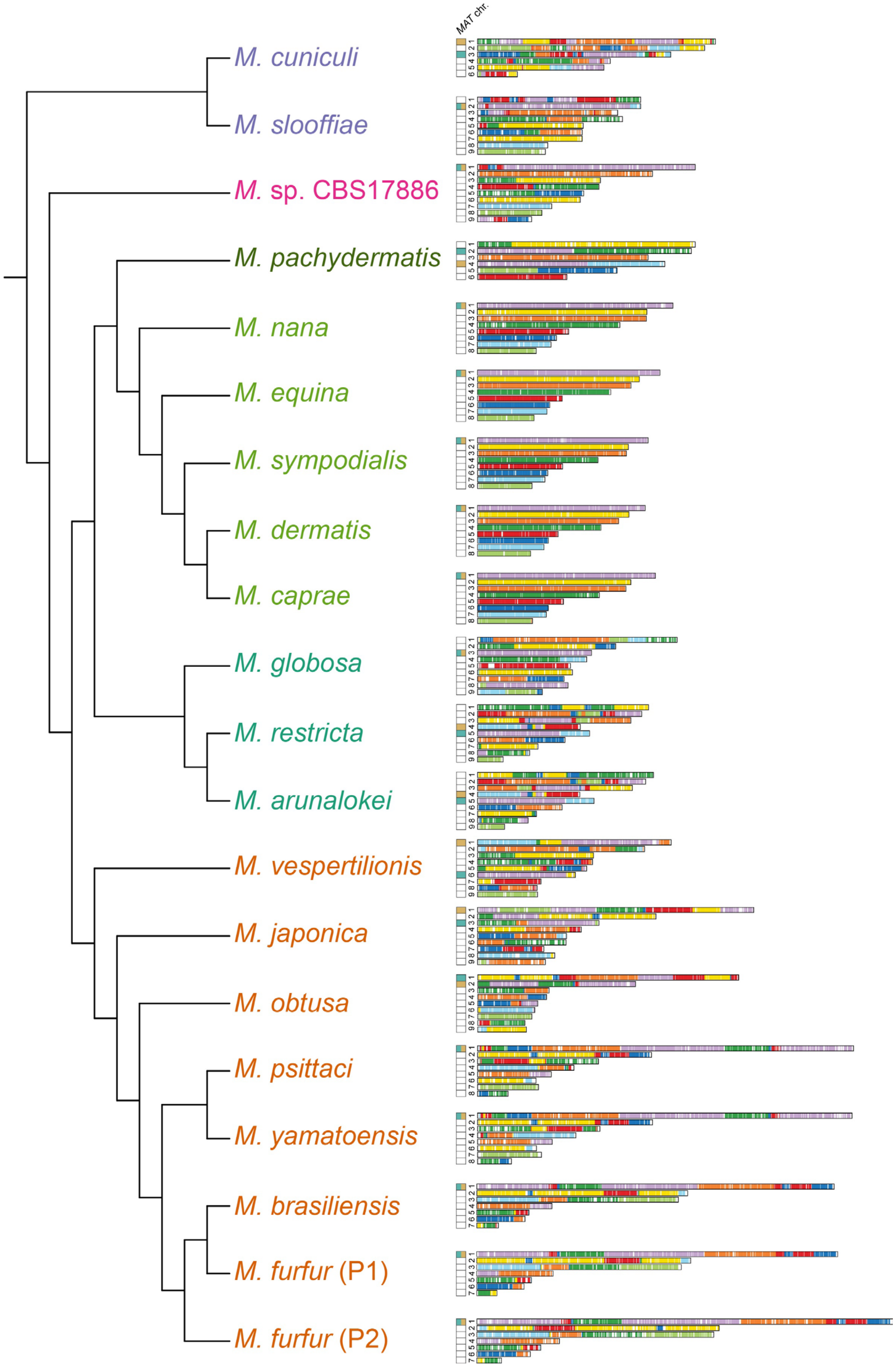


**Fig. S7.** Reconstruction of synteny blocks across *Malassezia* genomes. Conserved synteny blocks were identified between pairwise combinations of genomes with SynChro and were color-coded according to *M. sympodialis* chromosomes. Chromosome(s) containing the *P/R* (colored in teal) and *HD* (colored in gold) *MAT* loci are indicated for each species.

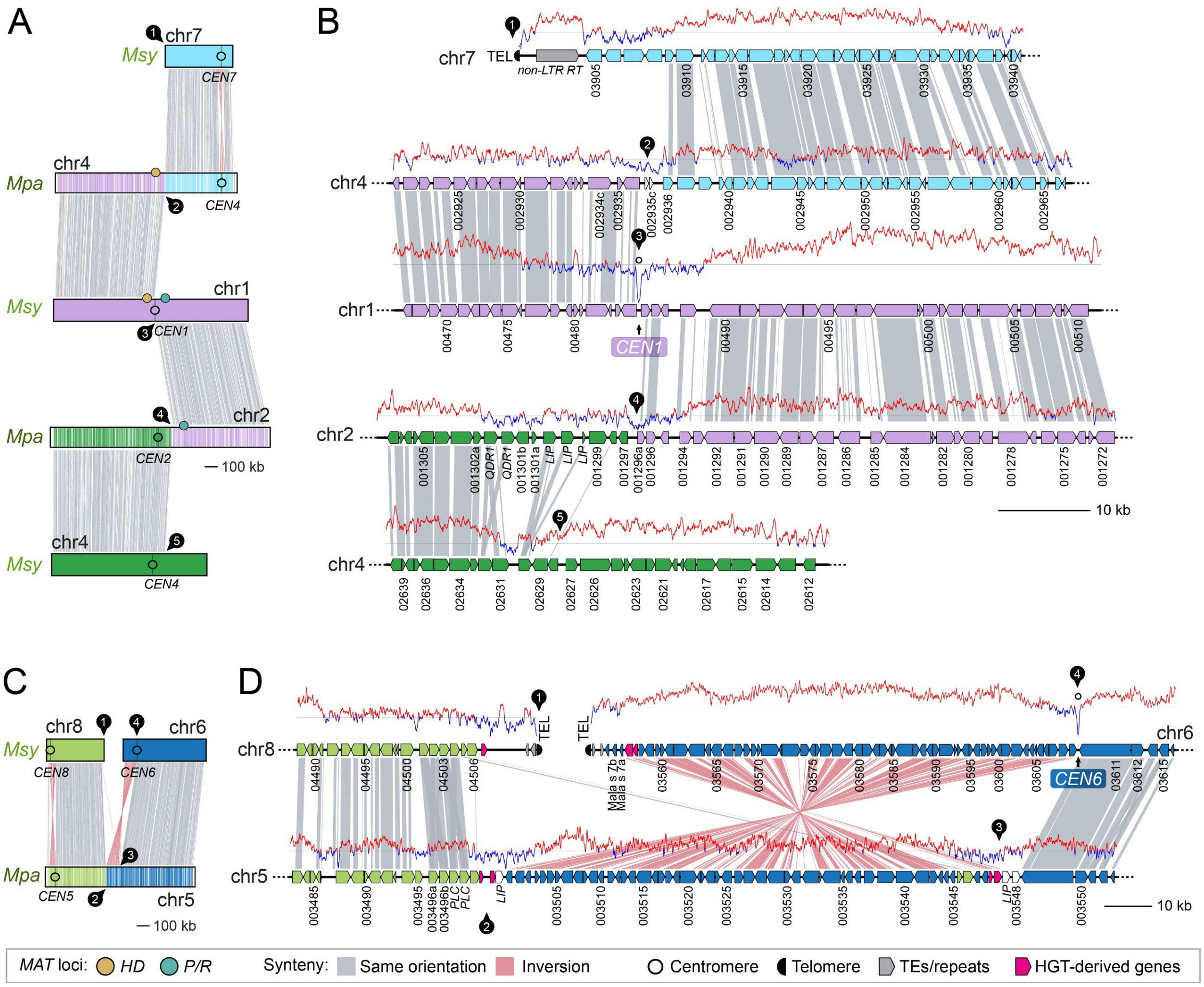


**Fig. S8.** Pseudobipolar-to-tetrapolar transition in *M. pachydermatis* and karyotypic reduction. (A) Synteny analysis comparing the *MAT* chromosome of *M. sympodialis* with corresponding regions in *M. pachydermatis*. The transition to tetrapolarity likely involved a centromere break followed by fusion of the acentric chromosome arms. This event is also associated with chromosome number reduction. Black circular labels mark the chromosomal regions associated with breaks or fusions, which are shown as zoomed-in views in panel (B). (C) Further chromosome number reduction in *M. pachydermatis* resulted from fusion of two chromosomes (chrs. 6 and 8) extant in *M. sympodialis* and in other species of clade B2. Synteny analysis allow us to hypothesize that a large inversion may have inactivated one of the centromeres (*CEN6*) after the fusion, possibly stabilizing the resulting chromosome. In panels (B) and (D), the GC content is shown as red/blue lines above each chromosomal region, with the grey line representing the average.

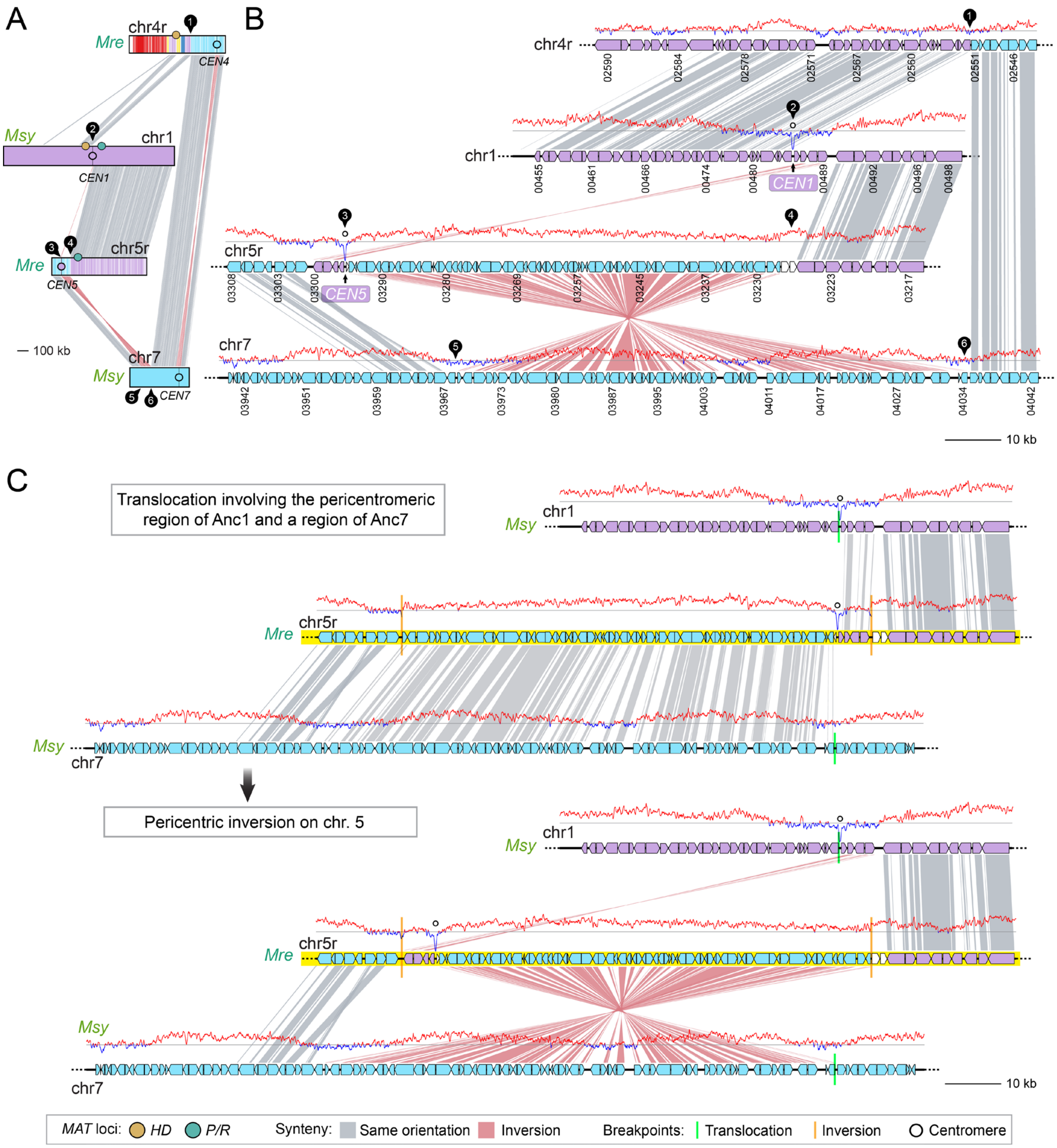


**Fig. S9.** Pseudobipolar-to-tetrapolar transition in *M. restricta* involved a translocation at the centromere-flanking region. (A and B) Synteny analysis comparing the *MAT* chromosome of *M. sympodialis* with corresponding regions in *M. restricta*. Black circular labels mark the chromosomal regions associated with rearrangements, which are shown as zoomed-in views in panel B. (C) Hypothesized rearrangements leading to tetrapolarity and extant chromosome configurations in *M. restricta*. First, a translocation within the Anc1 centromere-flanking region and a non-centromeric region of Anc7 relocated the two *MAT* loci onto different chromosomes. Second, a pericentric inversion in the derived *P/R*-containing chromosome relocated the Anc1 centromere (matching *M. restricta CEN5*) and a few flanking genes. The GC content is shown as red/blue lines above each chromosomal region, with the grey line representing the average.

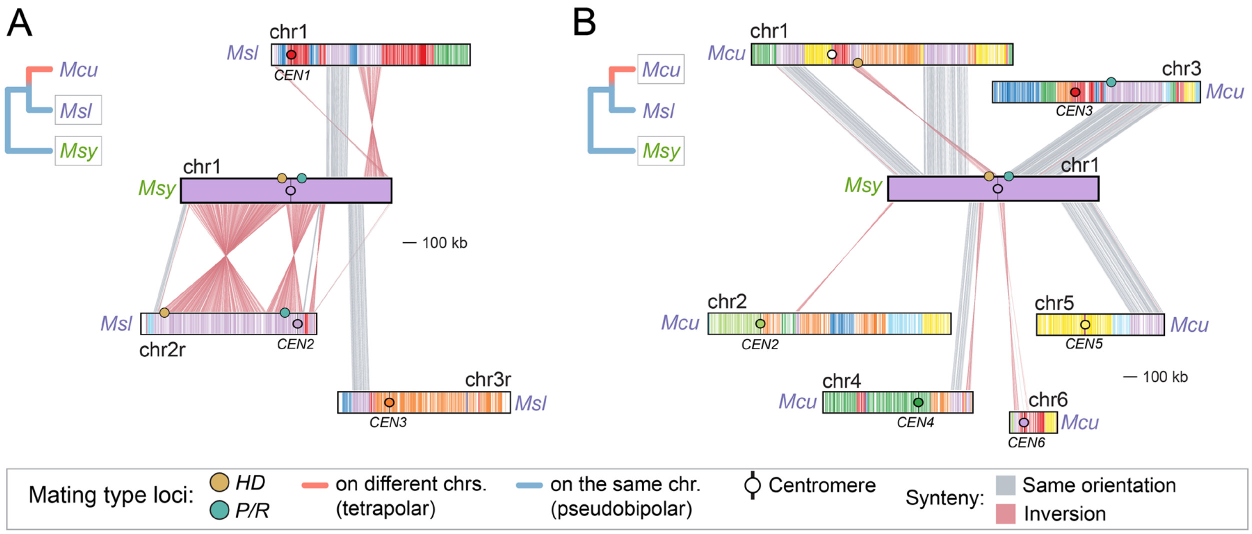


**Fig. S10.** The transition to tetrapolarity in *M. cuniculi* is complex. In this species the transition was associated with an intricate process of karyotypic changes and chromosome number reduction. Synteny comparison between the *MAT*-chromosome of *M. sympodialis* with corresponding regions in (A) *M. slooffiae* and (B) *M. cuniculi*. Note that the *M. sympodialis MAT*-chromosome is dispersed over the six chromosomes in *M. cuniculi* as opposed to only three in *M. slooffiae*.

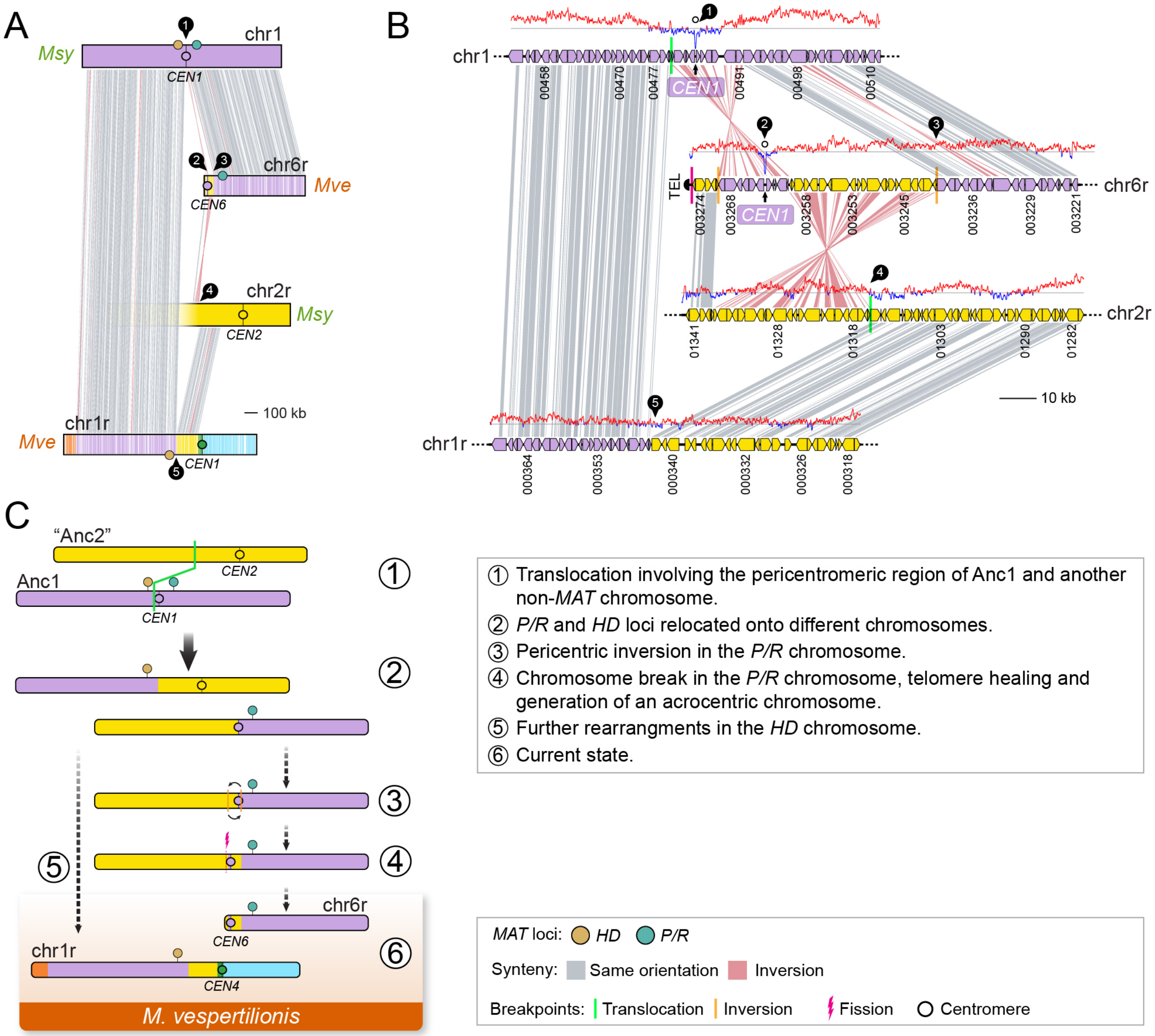


**Fig. S11.** Pseudobipolar-to-tetrapolar transition in *M. vespertilionis* involved a translocation at the centromere-flanking region. (A and B) Synteny analysis comparing the *MAT* chromosome of *M. sympodialis* with corresponding regions in *M. vespertilionis*. Black circular labels mark the chromosomal regions associated with rearrangements, which are shown as zoomed-in views in panel B. The GC content is shown as red/blue lines above each chromosomal region, with the grey line representing the average. (C) Proposed chromosomal rearrangements underlying the transition to a tetrapolar configuration from an ancestral pseudobipolar state.

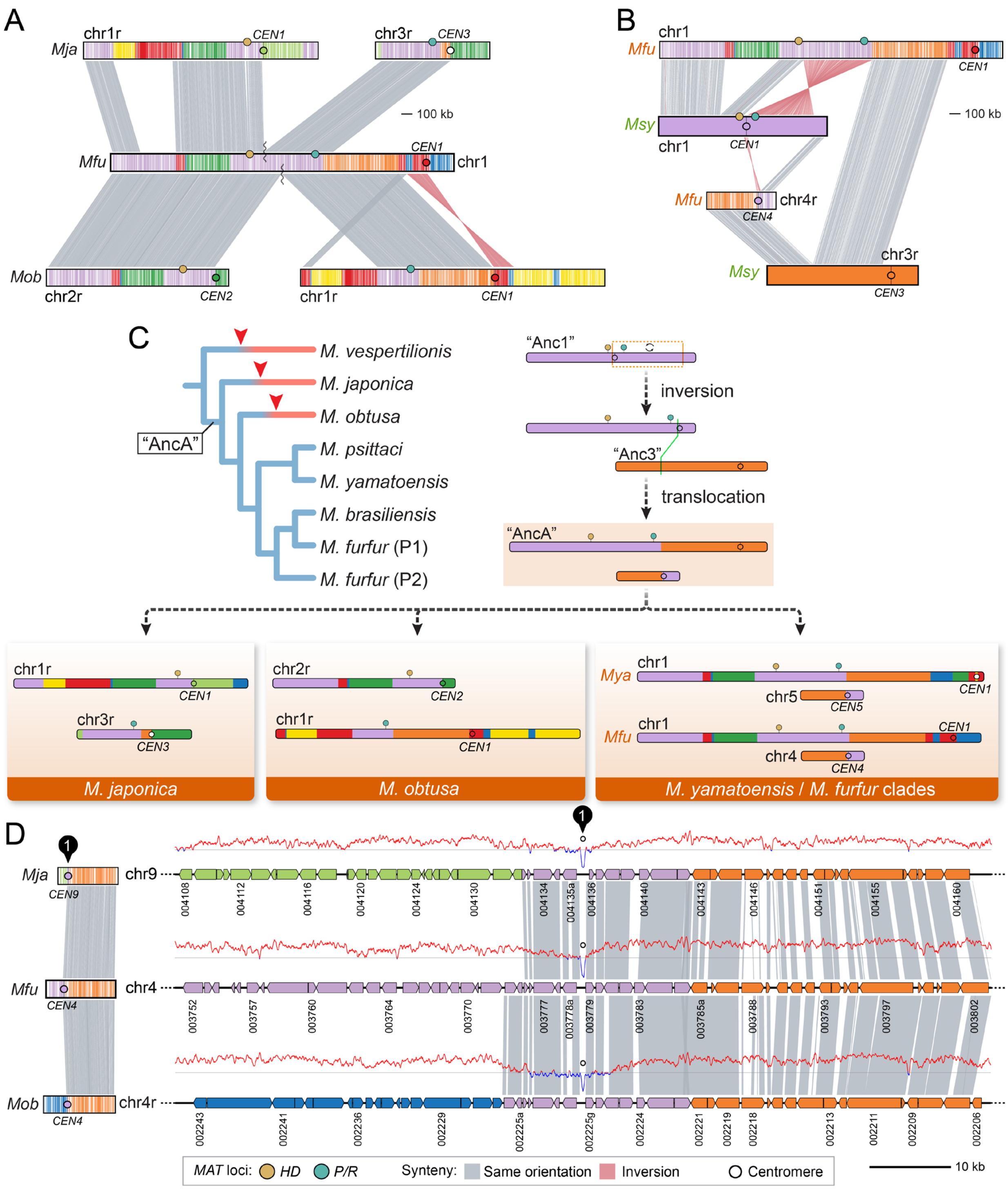


**Fig. S12.** Pseudobipolar to tetrapolar transition in *M. japonica* and *M. obtusa* and maintenance of a pseudobipolar structure in *M. furfur*/*M. yamatoensis* clades.

**Figure S12.** (continued). Synteny comparison between the *MAT* chromosome of *M. furfur* and corresponding regions in (A) *M. japonica* and *M. obtusa*, and in (B) *M. sympodialis*. (C) Proposed events underlying the formation of a modified pseudobipolar *MAT* chromosome (“AncA”) that conceivably emerged after the split from the *M. vespertilionisis* common ancestor. First, an inversion relocated the *P/R* locus farther apart from the *HD* locus and moved the centromere closer to the end of the chromosome. Subsequently, a translocation involving the inversion-derived chromosome and the “Anc3” displaced the “Anc1” centromere onto a different genomic context. The “AncA” chromosome underwent further species-specific rearrangements. In *M. japonica* and *M. obtusa* independently, one such rearrangement separated the *P/R* and *HD* loci onto different chromosomes. (D) Synteny comparison of extant chromosomes of *M. japonica*, *M. furfur*, and *M. obtusa* harboring the “Anc1” centromere. The GC content is shown as red/blue lines above each chromosomal region, with the grey line representing the average.

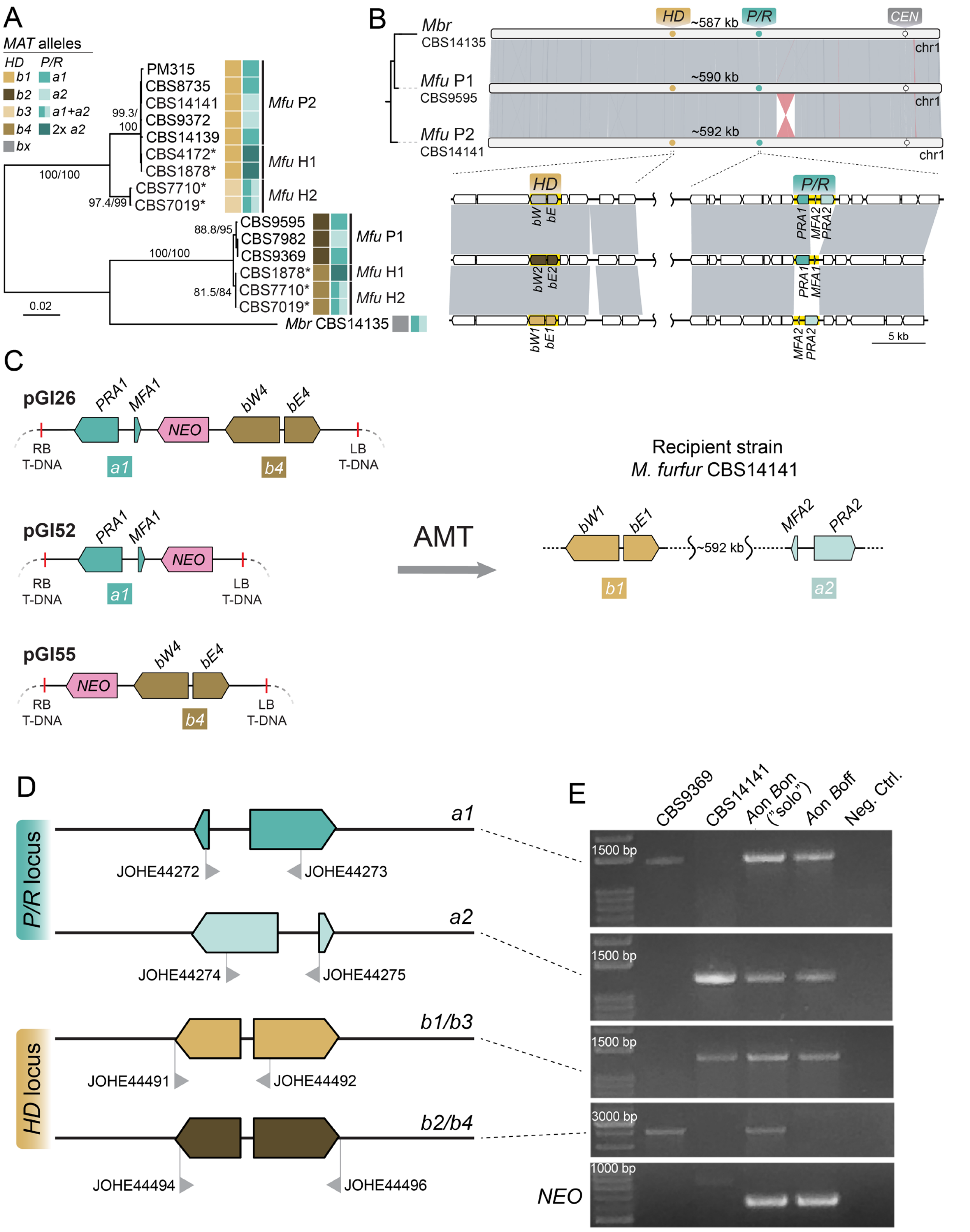


**Fig. S13.** *Malassezia furfur* strains and molecular confirmation of *A*on *B*off and *A*on *B*on ("solo") strains derived from *M. furfur* CBS14141.

**Fig. S13**. (continued). (A) Maximum likelihood phylogeny of *M. furfur* and *M. brasiliensis HD* alleles, and mating-type identity of the strains. The tree was constructed with IQ-TREE2 (model K3Pu+F+I) using 2,499 nucleotide sites. *M. furfur* hybrid strains (lineages H1 and H2) previously characterized (35) are marked with asterisks. (B) Synteny analysis along the *MAT* loci-containing chromosome of two *M. furfur* strains (P1 and P2) and *M. brasiliensis*. (C) Schematic representations of the T-DNA regions of: (i) plasmid pGI26 that contains the *a1-NEO-b4* transgene (upper panel); (ii) plasmid pGI52 harboring the *a1-NEO* transgene (middle panel); and (iii) plasmid pGI55 carrying the *NEO-b4* transgene (lower panel). These transgenes were inserted in the recipient strain *M. furfur* CBS14141 (*MAT a2b1*) through *A. tumefaciens* mediated transformation (AMT). RB and LB indicate the right and left borders of the T-DNA, respectively. (D) Schematic representations of *M. furfur P/R* and *HD* loci showing the positions of the primers specifically designed to detect the *a1*, *a2*, *b1/b3* and *b2/b4* alleles (35). (E) PCR screening of the *P/R* and *HD* alleles, and the *NEO* gene used as selectable marker. Note that strain CBS9369 (*MAT a1b2*) was the donor of the *P/R a1* allele to be transformed into *M. furfur* CBS14141, while the *HD b4* allele was obtained from the diploid hybrid strain CBS7710 (5, 35); CBS7710 was not included in the PCR screening because the presence of the exogenous *b4* allele in the *A*on *B*on ("solo") strain could be verified with the primers JOHE44494-JOHE44496, designed to amplify both the *b*2 and *b4* alleles (35).

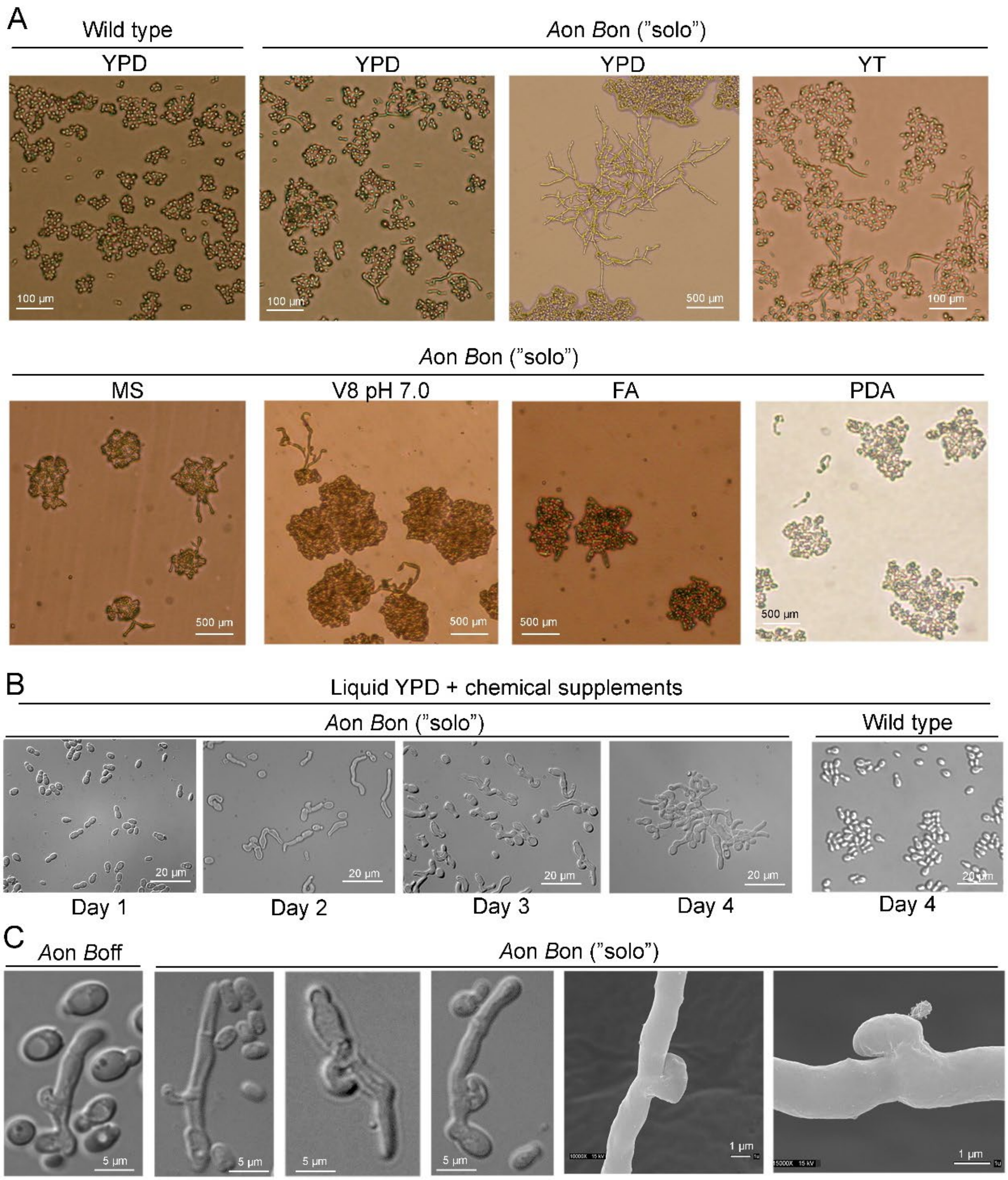


**Fig. S14.** Phenotypic features of the self-filamentous *A*on *B*on (“solo”) and *A*on strains derived from *M. furfur* CBS14141. (A) The *M. furfur A*on *B*on (“solo”) strain and the *M. furfur* CBS14141 wild type were streaked onto several agar media and incubated for 10 days at 32°C. Plates were directly observed by microscopy and photographed. MS is Murashige and Skoog medium; FA is filamentation agar; and PDA is potato dextrose agar. The strongest filamentation was triggered by YPD agar medium, and for this reason it was used to show a representative image of the morphology of the *M. furfur* CBS14141 wild type strain.

**Fig. S14**. (continued). (B) The *M. furfur Aon Bon* ("solo") and the *M. furfur* CBS14141 wild type strains were grown at 32°C in liquid YPD + supplements [squalene (1ml/L), potassium nitrate (1 g/L), sodium chloride (1.3 g/L), ferrous sulphate (0.01 g/L), magnesium sulphate (0.15 g/L) and glycine (3.75 g/L)], and an aliquot was observed on a daily basis by microscopy. A representative image of the *M. furfur* CBS14141 wild type is shown at the fourth day of incubation. (C) Magnifications of the clamp-like cells in the *A*on *B*off and *A*on *B*on ("solo") strains observed by optical microscopy or scanning electron microscopy (SEM) (last two panels on the right). Independent transformants obtained using the same transgenes also showed robust hyphal growth with septa and clamp-like connections.

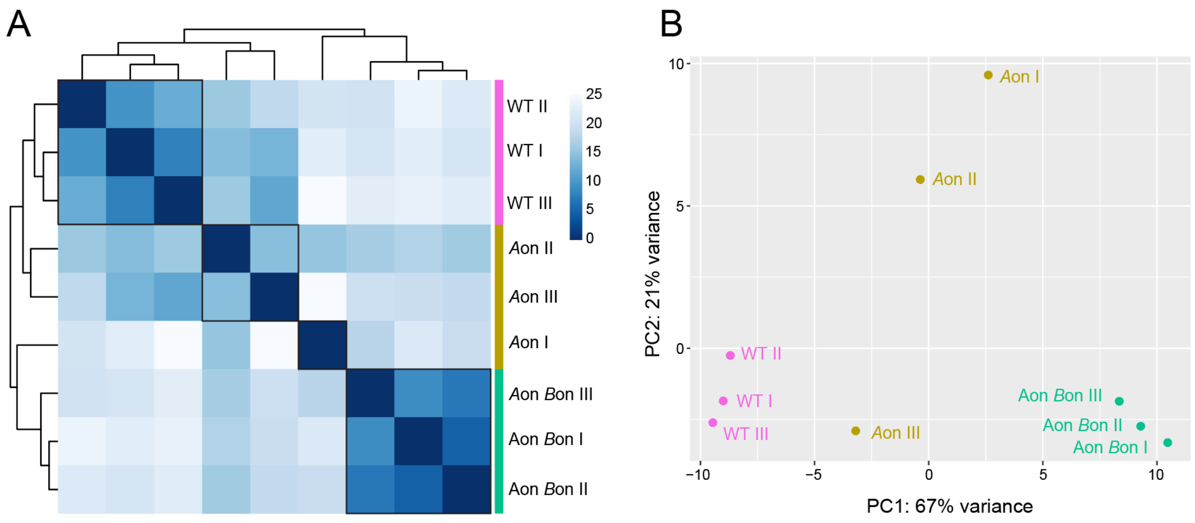


**Fig. S15.** (A) Heatmap reporting hierarchical clustering of gene expression level profiles of the different *M. furfur* strains (WT, *A*on and *A*on *B*on) upon regularized-logarithm (rlog) transformation of DESeq2-normalized expression data. Color codes (from white to dark blue) report Euclidean distances among samples, with dark blue corresponding to the maximum of correlation values (i.e., lower distances). (B) Principal component analysis (PCA) of the various samples, recapitulating observations from hierarchical clustering. Dots of the same color represent different replicates (I, II, and III) of the same sample. In both analysis we found some intrinsic variation among the *A*on replicates but removing sample '*A*on I' that clustered separately did not change the final set of differentially expressed genes obtained. These differences may be due to intrinsic variability or result from different proportions of yeast:hyphae cells at the end of the incubation period prior to isolation of RNA.

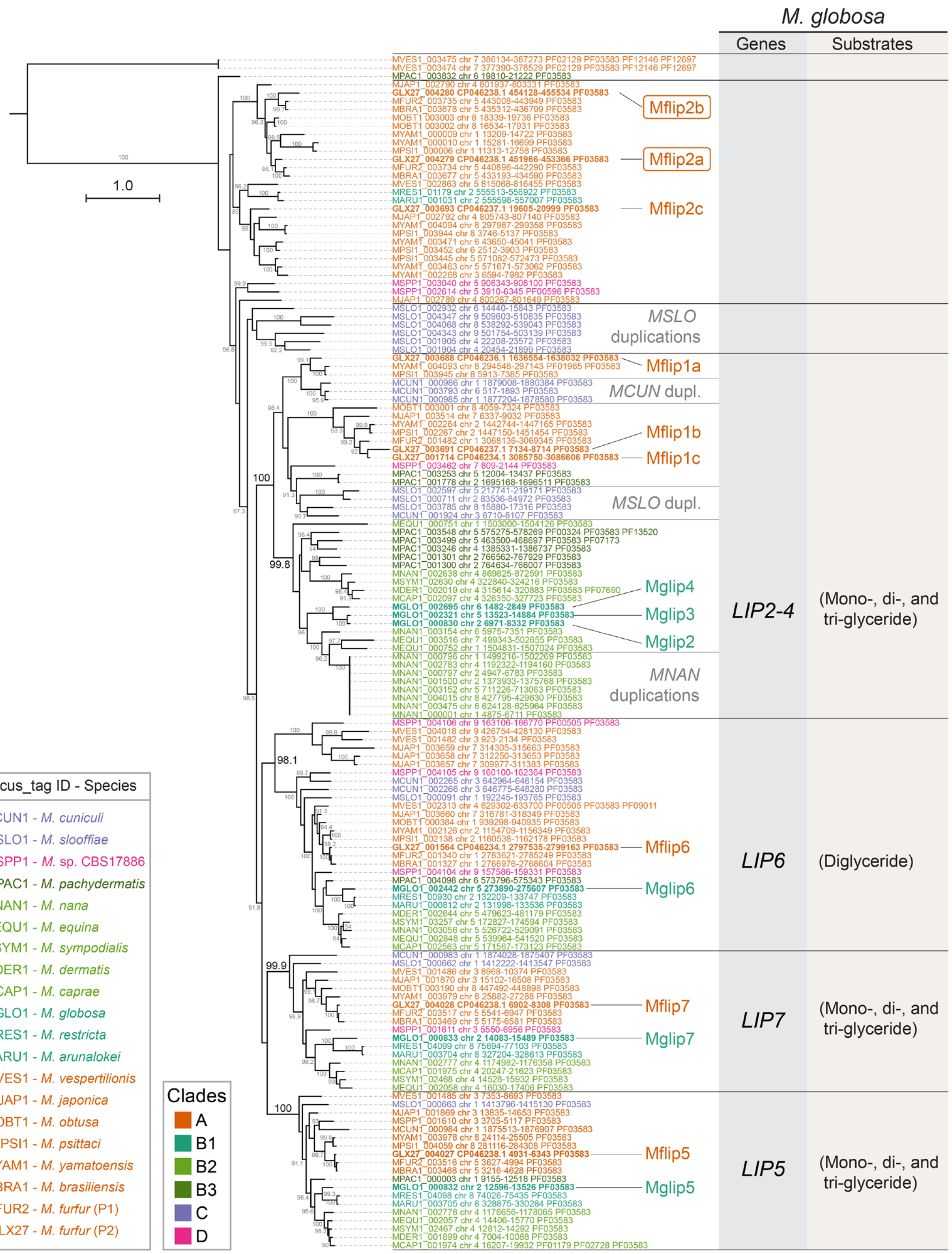


**Fig. S16.** ML phylogeny of predicted *Malassezia* secreted lipases (PF03583). Gene number expansion in each species seems to be largely due to recent species-specific duplications as seen in *M. slooffiae*, *M. nana*, and *M. globosa* (e.g., Mglip2, Mglip3 and Mglip4). Some of the lipases were characterized in previous studies by assessing their hydrolysis activity against various substrates upon expression in a heterologous system and are summarized here for *M. globosa* [reviewed in (43)]. *M. furfur MFLIP2a* and *MFLIP2b* genes are strongly upregulated in the *A*on *B*off and "solo" strains. The tree was inferred with IQ-TREE2 (model LG+F+I+R6) using 144 aligned and trimmed sequences identified by OrthoFinder. Branch lengths are given in number of substitutions per site (scale bar). Branch support (> 80%) is shown at the tree branches from 1,000 ultrafast bootstrap (UFboot) replicates. The tree is rooted at the midpoint.

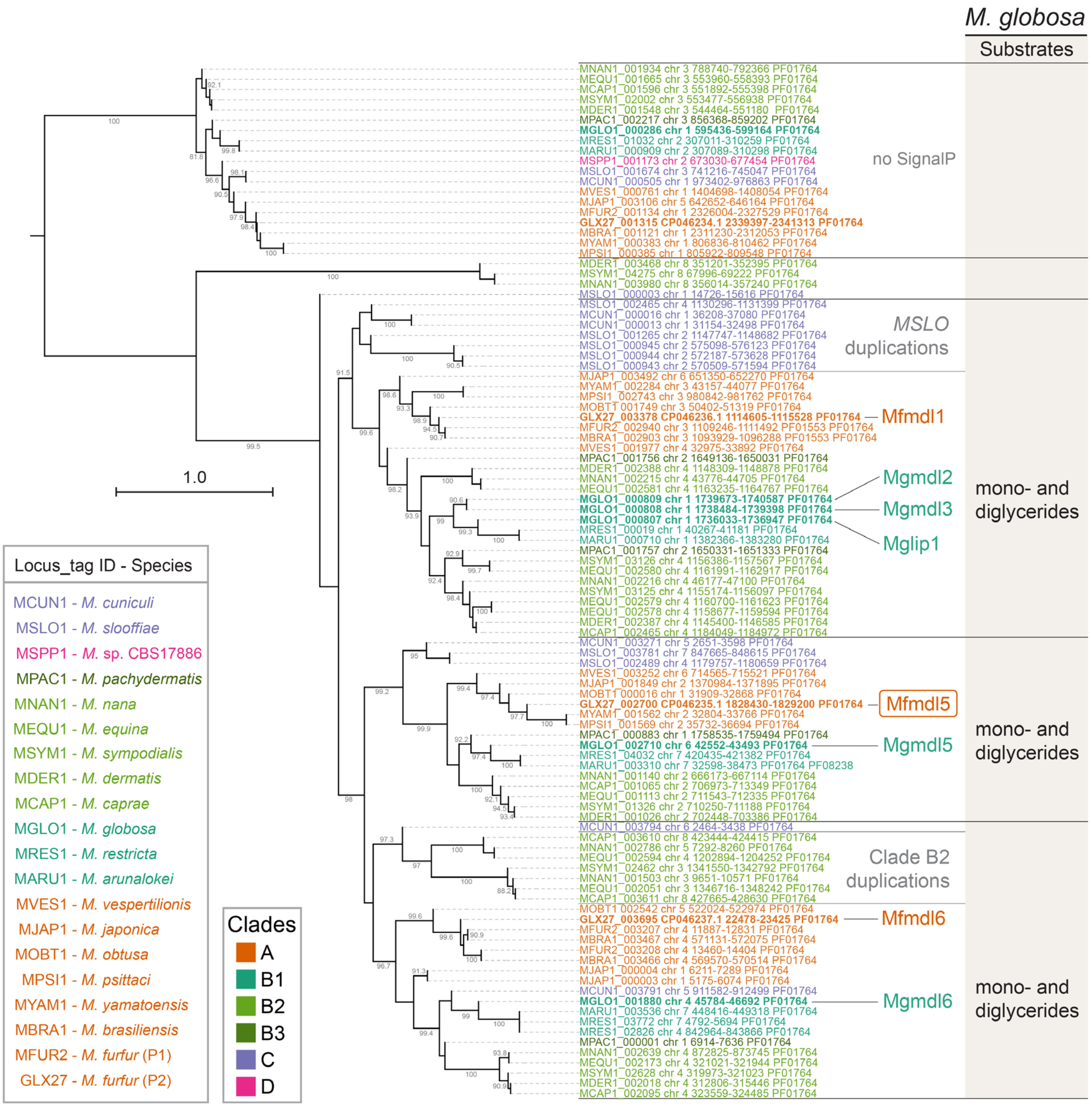


**Fig. S17.** ML phylogeny of predicted *Malassezia* Class 3 family lipases (PF01764). Some of these lipases were characterized in previous studies by assessing their hydrolysis activity against various substrates upon expression in a heterologous system and are summarized here for *M. globosa* [reviewed in (43)]. The *M. furfur* gene *MFMDL5* is among the upregulated genes in the *A*on *B*off and “solo” strains. The tree was constructed with IQ-TREE2 (model WAG+I+G4) using 101 aligned and trimmed sequences identified by OrthoFinder. Branch lengths are given in number of substitutions per site (scale bar). Branch support (> 80%) is shown at the tree branches from 1,000 ultrafast bootstrap (UFboot) replicates. The tree is rooted at the midpoint.

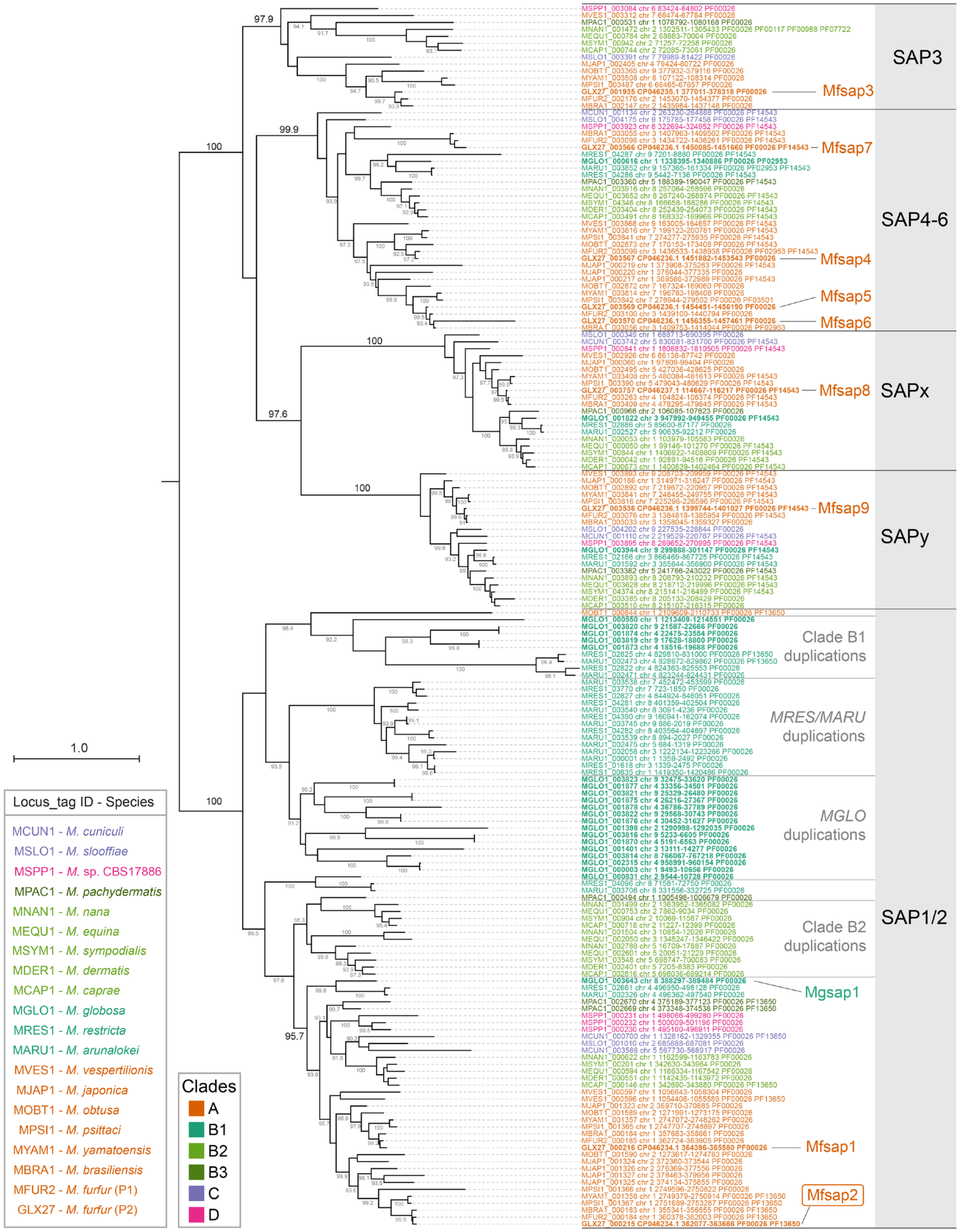


**Fig. S18.** ML phylogeny of predicted *Malassezia* secretory aspartyl proteases (PF00026). The *M. furfur MFSAP2* gene expression level in the WT is residual but it is strongly upregulated in the *A*on *B*off and "solo" strains. The tree was constructed with IQ-TREE2 (model LG+F+I+R6) using 176 aligned and trimmed sequences identified by OrthoFinder. Branch lengths are given in number of substitutions per site (scale bar). Branch support (> 80%) is shown at the tree branches from 1,000 ultrafast bootstrap (UFboot) replicates. The tree is rooted at the midpoint.

**Table S1.** Summary of genome assembly statistics and other genomic features.

| Species | Isolate | Assembly Size (bp)[a] | Number of | | | | | Telomeric repeats | |
|---|---|---|---|---|---|---|---|---|---|
| | | | ctgs[c] | % GC | Genes | Proteins | tRNAs | Left end | Right end |
| *M. cuniculi* | CBS11721 | 7,453,460 | 6 | 59.17 | 3,953 | 3,872 | 81 | TAACCC | GGGTTA |
| *M. slooffiae* | CBS7956 | 8,846,367 | 9 | 66.33 | 4,358 | 4,303 | 55 | TAACCC | GGGTTA |
| *Malassezia* sp. | CBS17886 (=46375-004) | 9,312,381 | 9 | 66.74 | 4,251 | 4,191 | 60 | TAACCC | GGGTTA |
| *M. pachydermatis* | KCTC27587 | 8,283,742 | 6 | 55.21 | 4,199 | 4,122 | 77 | TAACAC | GAGTTA |
| *M. nana* | CBS9557 | 7,702,145 | 8 | 57.97 | 4,032 | 3,959 | 73 | TTAACAC | GTGTTAA |
| *M. equina* | CBS12830 | 7,775,479 | 8 | 58.06 | 3,750 | 3,686 | 64 | TTAACAC | GTGTTAA |
| *M. sympodialis* | ATCC42132 | 7,747,459 | 8 | 59.07 | 4,588 | 4,484 | 77 | TTAACAC | GTGTTAA |
| *M. dermatis* | JCM11348 | 7,547,282 | 8 | 58.99 | 3,502 | 3,448 | 54 | n.d. | |
| *M. caprae* | CBS10434 | 7,728,824 | 8 | 59.83 | 3,629 | 3,561 | 68 | TTAACAC | GTGTTAA |
| *M. globosa* | CBS7966 | 9,084,671 | 9 | 52.06 | 4,081 | 4,000 | 81 | TAACAC | GTGTTA |
| *M. restricta* | KCTC27527 | 7,330,907 | 9 | 55.79 | 4,493 | 4,390 | 74 | TAACAC | GTGTTA |
| *M. arunalokei* | CBS13387 | 7,331,251 | 9 | 55.59 | 3,858 | 3,786 | 72 | TAACAC | GTGTTA |
| *M. vespertilionis* | CBS15041 | 7,595,203 | 9 | 56.64 | 4,022 | 3,964 | 58 | TAACCC | GGGTTA |
| *M. japonica* | CBS9431 | 8,433,779 | 9 | 62.29 | 4,373 | 4,315 | 58 | TAATCC | GGATAA |
| *M. obtusa* | CBS7876 | 8,169,464 | 9 | 62.98 | 3,400 | 3,338 | 62 | TAATCC | GGATAA |
| *M. psittaci* | CBS14136 | 8,186,253 | 8 | 49.71 | 4,064 | 4,006 | 58 | TAATCC | GGATAA |
| *M. yamatoensis* | CBS9725 | 8,264,016 | 8 | 49.64 | 4,103 | 4,047 | 56 | TAATCC | GGATAA |
| *M. brasiliensis* | CBS14135 | 8,000,312 | 7 | 63.17 | 3,967 | 3,911 | 56 | TAATCC | GGATAA |
| *M. furfur* P1 | CBS9595 | 8,119,594 | 7 | 64.49 | 4,011 | 3,951 | 60 | TAATCC | GGATAA |
| *M. furfur* P2 | CBS14141 | 8,268,083 | 7 | 64.9 | 4,601 | 4,533 | 68 | TAATCC | GGATAA |

Genomes produced in this study are marked in grey.
[a] nuclear genome size only; the mitochondrial genome was not assembled in this study.
[b] telomeres not detected (n.d.) in the available assembly.
[c] the number of contigs (ctgs) is equivalent to the number of chromosomes. *M. dermatis*, was not re-sequenced in this study and the available genome still has gaps and lack telomeric repeats. All other genomes are assembled telomere-to-telomere and do not contain any gap. The rDNA cluster is assembled as a collapsed region (i.e., only contain a few copies of the array).

**Table S2.** *M. furfur* and *M. sympodialis* strains used in fusion and mating assays, and *M. furfur* transformants obtained in this study.

| Species | strain | Remarks | Mating type[a] | Reference |
|---|---|---|---|---|
| *M. furfur* | CBS9595 | P1 lineage | *a1b2* | (35) |
| *M. furfur* | CBS9575 | P1 lineage | *a2b2* | (35) |
| *M. furfur* | CBS9574 | P1 lineage | *a1b2* | (35) |
| *M. furfur* | CBS9589 | P1 lineage | *a1b2* | (35) |
| *M. furfur* | CBS7982 | P1 lineage | *a2b2* | (35) |
| *M. furfur* | CD866 | P1 lineage | *a2b2* | (35) |
| *M. furfur* | CBS9369 | P1 lineage | *a1b2* | (35) |
| *M. furfur* | CBS8735 | P2 lineage | *a1b1* | (35) |
| *M. furfur* | CBS14139 | P2 lineage | *a1b1* | (35) |
| *M. furfur* | CBS14141 | P2 lineage | *a2b1* | (35) |
| *M. furfur* | CBS14141 *ade*2Δ*NAT* | P2 lineage | *a2b1* | (2) |
| *M. furfur* | PM315 | P2 lineage | *a1b1* | (35) |
| *M. furfur* | CBS7710 | H2 lineage | *a1a2b3b4* | (5,35) |
| *M. sympodialis* | ATCC42132 *NAT* | *NAT* resistant | *a1b2* | (2) |
| *M. sympodialis* | ATCC44340 *NEO* | *NEO* resistant | *a2b2* | (2) |
| *M. furfur* | *A*on *B*off | *NEO* resistant | *a1a2b1* | This study |
| *M. furfur* | *A*on *B*on (“solo”) | *NEO* resistant | *a1a2b1b4* | This study |

[a]mating type were designated based on PCR amplification of the *P/R* (*a*) locus and PCR amplification plus sequencing of the *HD* (*b*) locus.

**Table S3.** List of primers obtained in this study.

| Primer ID | Sequence (5' > 3') | Purpose |
|---|---|---|
| JOHE43874 | GCGCGCCTAGGCCTCTGCAGGTCGACTCTTTG ACTTGTGACTTGAATGC | *MAT a1* cloning - F |
| JOHE43875 | GAGGATCTGCACCGTGGAATCCTGCCTCTTCT AGCTTGC | *MAT a1* cloning - R |
| ALID2078 | TCCACGGTGCAGATCCTC | *NEO* marker amplification - F |
| ALID2081 | CGTCCTCTCCTATGTCTG | *NEO* marker amplification - R |
| JOHE43876 | CAGACATAGGAGAGGACGGCTAACCATCGCC GCGATCC | *MAT* b4 cloning - F |
| JOHE43877 | TGATTACGAATTCTTAATTAAGATATCGAGAGT TTGGCACCCGTCAGACG | *MAT* b4 cloning - R |
| JOHE43279 | CT*GATC*CAAGCTCAAGCTC | Assess correct recombination with pGI3 at the 5' junction - F |
| JOHE43281 | GTCGGAGAAGCAGTCAATGC | Assess correct recombination with pGI3 at the 5' junction - R |
| JOHE43280 | GTTGGCCGATTCATTAATGC | Assess correct recombination with pGI3 at the 3' junction - F |
| JOHE43282 | CACCAGGGTTTCCAGTCTC | Assess correct recombination with pGI3 at the 3' junction - R |
| JOHE44551 | TGATTACGAATTCTTAATTAAGATATCGAGCGT CCTCTCCTATGTCTG | recombination *NEO* with pGI3 at the 3' - R |
| JOHE46235 | GCGCGCCTAGGCCTCTGCAGGTCGACTCTTCC ACGGTGCAGATCCTCGG | recombination *NEO* with pGI3 at the 5' - F |
| JOHE44272 | AACCATCCATGCTGACATTT | *M. furfur MAT a1* -F |
| JOHE44273 | TTGGCAGAGTTGACAGGCT | *M. furfur MAT a1* -R |
| JOHE44274 | GAGCCACAAGATAATGTCAA | *M. furfur MAT a2* -F |
| JOHE44275 | AGACTTCCTGAACAGTGTCC | *M. furfur MAT a2* -R |
| JOHE44491 | TTCGGTTGACGGTCCCTCGGC | *M. furfur MAT b1-b3* - F |
| JOHE44492 | ACCGCGACTGCGCATCCGCG | *M. furfur MAT b1-b3* - R |
| JOHE44494 | TTCGCCAAATGTGTTCGGCC | *M. furfur MAT b2-b4* - F |
| JOHE44496 | CAGCAACACCCGCCTCGCTT | *M. furfur MAT b2-b4* - R |
| JOHE43965 | GCTTGGGTGGAGAGGCTATT | internal *NEO* F |
| JOHE43966 | AGCCAACGCTATGTCCTGAT | internal *NEO* R |

**Dataset S1 (separate file).** Genome assembly, genomic features and information on raw sequencing data generated in this study. (A) Genome sequencing, assembly, and polishing approaches. (B) NCBI accession numbers of each genome and raw read data generated and used in this study. (C) Predicted centromere locations of the genome assemblies generated in this study. (D) Predicted non-long-terminal repeat (non-LTR) sequences identified at telomere-adjacent regions (only the product of the longest open reading frame (ORF) is given for each species). (E) Mating type (*MAT*) loci chromosomal locations in *Malassezia*.

**Dataset S2 (separate file).** List of upregulated and downregulated genes between (A) *A*on *B*off vs. Wild-type (WT) and (B) *A*on *B*on ("solo") vs. WT using DESeq2. logFC = log2 fold-change; lfcSE = log fold change Standard Error; padj = false discovery rate (FDR) adjusted for multiple testing with the Benjamini-Hochberg procedure.

**Dataset S3 (separate file).** DESeq2 normalized read counts (median of ratios method of normalization) for all genes and samples.

**Dataset S4 (separate file).** Differential expression analysis (all genes) comparing (A) *A*on *B*off vs. Wild-type (WT) and (B) *A*on *B*on ("solo") vs. WT using DESeq2. logFC = log2 fold-change; lfcSE = log fold change Standard Error; padj = false discovery rate (FDR) adjusted for multiple testing with the Benjamini-Hochberg procedure.

**Disclaimer**

Any use of trade, firm, or product names is for descriptive purposes only and does not imply endorsement by the U.S. Government.
